# Developmental vitamin A deficiency induces sex-specific reward processing alterations through a dysregulation of the mesolimbic dopamine transmission in mice

**DOI:** 10.1101/2025.08.28.672827

**Authors:** Pauline Couty, Sonya Yung, Ilona Dulapt, Loreleï Berger, Adrien Santoro, Lola Hardt, Anna Petitbon, Fabien Ducrocq, Roman Walle, Julien Catanese, Serge Alfos, Jean-Christophe Helbling, Maria-Florencia Angelo, Charlotte Sabran, Patrick Borel, Guillaume Ferreira, Clémentine Bosch-Bouju, Pierre Trifilieff, Katia Touyarot

## Abstract

Neurodevelopmental psychiatric diseases such as schizophrenia or affective disorders share common symptomatic dimensions, in particular reward processing dysfunctions, associated with dysregulation of dopamine (DA) transmission. Retinoic acid (RA) homeostasis is altered across psychiatric disorders but whether impaired developmental RA signaling impacts the functionality of DA-related reward processing at adulthood remains poorly explored. Herein, we explored in male and female mice how developmental vitamin A deficiency (VAD), as a model of blunted RA signaling, could impact motivational processes through a modulation of mesolimbic DA transmission. Behavioral performances were evaluated using operant conditioning tasks, parallel with investigations of the integrity of DA transmission through biochemical analyses of markers of DA transmission and measures of DA dynamics using DA biosensor coupled with fiber photometry. Finally, chemogenetic manipulation of the mesolimbic DA pathway was used to normalize DA transmission and assess the effect on motivational performance in VAD offspring. Developmental VAD induced sex-specific alterations of reward-related processes at adulthood. Indeed, while female behavioral performances were spared, VAD males exhibited elevated instrumental performance and impulsivity. These behavioral alterations were coherent with reduced DA transporter (DAT) expression and increased DA dynamic in the mesolimbic pathway. Strikingly, chemogenetic inhibition of the mesolimbic DA pathway normalized motivational performance in VAD males. Our results show that developmental RA hyposignaling induces sex-specific reward processing alterations in adulthood through hyperactivity of the mesolimbic DA pathway. Our data support that developmental impairment in RA signaling might be at the core of reward-related symptoms across psychiatric disorders.

## INTRODUCTION

Psychiatric conditions such as schizophrenia or affective disorders are multifactorial neurodevelopmental diseases that involve both genetic and environmental factors. While these diseases are classically considered clinically different, they display common symptoms that share similar etiology and neurobiological substrates. Specifically, alterations of reward processing (1–4), particularly motivational alteration and impulsivity, belong to a common symptomatic frame that correlates with a dysregulation of the mesolimbic dopamine (DA) transmission (1,5). However, whether such impairments relate to similar pathogenic mechanisms remains unclear.

In this context, the important role of retinoic acid (RA), the active metabolite of vitamin A, during brain development (6) led to the hypothesis that altered RA signaling could be involved in the etiology of neurodevelopmental psychiatric disorders (7), though direct evidence remains sparse. A large population-based birth cohort study reported that a low maternal retinol (i.e. vitamin A) serum concentration during pregnancy increased the risk for schizophrenia in offspring by more than threefold (8). In line with this, low plasma retinol or RA concentrations have been observed in patients with developmental psychiatric conditions such as schizophrenia or attention deficit hyperactivity disorder (9–13). Beyond changes in RA levels *per se*, various polymorphisms in genes involved in RA signaling, including RA nuclear receptors or RA synthesis enzymes, have been identified in schizophrenia and affective disorders, and altered gene expression of key mediators of RA signaling have also been found in postmortem brain tissue across these developmental psychiatric disorders (14–20). Of note, retinoid analogs have been proposed as candidates in the treatment of schizophrenia or autism (21–23). However, little mechanistic data link dysfunctions of developmental RA signaling with neurological and behavioral features typical of psychiatric disorders.

The implication of RA signaling for brain development has been largely demonstrated for the formation of the neural plate, but also at later stages for the differentiation of various neuronal types (6). Regarding DA pathways, while only indirect evidence suggest that RA promotes differentiation into dopaminergic neurons (24,25), it is well established that RA is involved in the development and differentiation of striatal dopaminoceptive neurons (26–28). RA mostly acts by regulating transcription of various genes containing regulatory sequences in their promoter through its action on two receptor types, RARs (retinoic acid receptor) and RXRs (retinoid x receptor) (29,30). Consistent with its enrichment in RA receptors, the striatum is highly vulnerable to altered developmental RA signaling (31,32). Indeed, genetic ablations of retinoid receptors or aldehyde dehydrogenase enzyme, whose main function is to synthesize RA from retinol, decrease DA receptors expression, particularly in the ventral striatum, i.e. the nucleus accumbens (NAc) (33–36). Functionally, those genetic ablations lead to abnormal striatal functions as revealed by motor and affective deficits at adulthood (33,35). Beyond developmental effects, viral-mediated manipulations of RA signaling in the NAc at adulthood can bidirectionally modulate reward processing, particularly motivational processes (37), while manipulation of retinoid receptors levels in the NAc impacts reward sensitivity, by altering the excitability of dopaminoceptive neurons (38). By contrast, vitamin A supplementation appears as a preventive strategy to improve striatal functions in a rat model of Parkinson disease by increasing striatal expression of retinoid and DA receptors (39). However, the developmental impact of low RA signaling on the integrity of DA transmission *per se and* associated behavioral dimensions remains poorly explored.

Here, we found that prenatal vitamin A deficiency (VAD), which induces RA hyposignaling in the mesolimbic pathway, leads to aberrant reward processing at adulthood in males but not in female mice. Such a sex effect relates with impaired expression of the DA transporter (DAT) selectively in males, that correlates with enhanced DA dynamics in the mesolimbic pathway. Strikingly, chemogenetically dampening DA transmission normalizes reward processing in VAD male mice.

## MATERIALS AND METHODS

### Animals and diets

Animals were housed in groups in standard cages with a 12h light/dark cycle, in an enriched and controlled environment (22°C ± 2°C, 40% of humidity), with food and water *ad libitum*. The following mouse lines were used in this study: C57BL6/J wild-type (WT) from Janvier Labs and DAT-Cre mice (DAT^IREScre^, B6.SJL-Slc6a3tm1.1(cre)Bkmn/J, RRID: IMSR_JAX:006660). Female WT were crossed with WT or heterozygous DAT-Cre males to generate WT or single transgenic DAT-Cre offsprings, respectively. Female breeders were given access to control diet (CD: 7.5 International Unit (IU, 1IU=0,3µg of retinol) of retinol per g of diet) or vitamin A deficient diet (VAD: 0.4 IU) across gestation and lactation, and offspring were maintained with the same diet, from weaning. The different diets were isocaloric and differed only by the amount of retinol. All experiments were performed according to the criteria of the European Communities Council Directive (2010/63/UE) and were approved by the local ethical committee and the French Ministry of Higher Education and Research (authorization 34547-2022010516039112).

### Stereotaxic surgery

Analgesia was induced with a subcutaneous injection of Buprenorphine (BUPRECARE, 0.1 mg/kg) 30 minutes prior to the surgical procedure. Then mice were anesthetized with isoflurane (induction at 5%, procedure at 1-2%) and then placed in a stereotactic frame (RWS Life Science) on a heating pad (37°C). AAV were injected using a 10µl Hamilton syringe (Hamilton). The syringe was left in place for 5 minutes post-injection to prevent backflow before being slowly removed. Postoperative analgesia was ensured with a subcutaneous injection of NSAID (Non-Steroidal Anti-Inflammatory Drug) (CARPROX, 5mg/kg). Mice were placed on a heating pad until full recovery from the procedure and were monitored for 3 days post-surgery. Behavioral experiments were performed 3 to 4 weeks after the surgeries.

For fiber photometry experiments, we unilaterally injected 400 nL of adeno-associated viral (AAV) vectors bearing the DA sensor dLight 1.3b (AAV9-hSyn1-chl-dLight1.3b-WPRE-bGHp(A), titer: 7.9 x 10^12^ vg/ml, ETH Zurich) into the NAc (AP=+1.7, ML=+/-1.10, DV=-4.4 mm) at a rate of 100 nL/min. The injected hemisphere (right or left) was assigned randomly for each mouse. The optic fiber (Fiber Optic Cannula with Ceramic Ferrule, 0.D. 1.25 mm, 05 NA, 400µm core diameter, L=6 mm; R-FOC-BL400C-50NA, RWD) was placed 250 µm above the injection site (DV=-4.15 mm) and immobilized with an opaque dental cement (Super-Bond, Sun Medical, Meliodent, Kulzer). For chemogenetic experiments, we bilaterally injected 300 nL of AAV bearing the inhibitory Designer-Receptor Exclusively Activated by Designer Drug (DREADD) HM4Di (AAV8-hSyn1-dlox-HM4D(Gi)-mcherry(rev)-dlox-WPRE-hGHp(A), titer: 6.3 x 10^12^ vg/ml or AAV8-hSyn1-dlox-mCherry(rev)-dlox-WPRE-hGHp(A), titer: 5.9 x 10^12^ vg/ml, ETH Zurich) into the ventral tegmental area (VTA) (AP=-3.3, ML=+/-0.50, DV=-4.30) at a rate of 50 nL/min. The skin was then sutured.

### Behavioral experiments

#### Operant chamber apparatus

The operant chambers (Imetronic, France) had internal dimensions 30x46x36 cm and were located in a soundproof and light-attenuating cabinet. Each operant chamber had two opaque panels at the right and left walls and two clear Plexiglas panels at the back and front walls and was illuminated throughout all sessions with a houselight located at the top of the chamber. The chambers were equipped with two retractable levers (2x4x1 cm). Rewarded lever presses were indicated by a brief tone (65 db, 3000 Hz, 200 ms) and a cue light, mounted 8,5 cm above the lever. A food port was positioned 4 cm above the grid floor at equidistance between the two levers. Dehydrated milk pellets without vitamin A were used as a reward (Rodent Pellet Tablet (20 mg) with non-added vitamin A, non-flavoured, 5BUP, Test Diet). For each mouse, all the operant procedures were executed in the same operant chamber. One session was run each day, 5 per week. Operant chambers were connected to a computer equipped with POLY Software (Imetronic), which allowed for control of the session parameters and recording of the data. Animals were food-restricted in order to maintain them at 85-90% of their *ad libitum* weight. For the study of motivational processes, only one of the two levers was reinforced, i.e., associated with the release of the reward. Pressing the unreinforced lever had no consequence. Mice were first trained in Pavlovian (Supplemental Methods) and instrumental conditioning before being subjected to motivational tasks.

#### Instrumental conditioning

Mice were trained to press on the reinforced lever (RL) to obtain a reward under a fixed ratio 1 (FR1) schedule of reinforcement, in which each lever press leads to reward delivery. The session began with the illumination of the house light and the insertion of the levers inside the chamber. The mice were exposed to this schedule until obtaining 40 rewards or after one hour, during approximately 5 sessions. Then the animals were shifted to a random ratio (RR) schedule of reinforcement, in which lever presses were reinforced with a given reward probability in order to increase their ability to press. The training consisted of three consecutive steps characterized by a different RR value that defined the probability of reinforcement: RR5 (p=0.2), RR10 (p=0.1), and RR20 (p=0.05). The mice were exposed to this schedule until obtaining 40 rewards or after 30 minutes, during approximately 3 or 4 sessions for each step.

#### Progressive ratio task

In the progressive ratio schedule (PRx2), the number of lever presses required to obtain a reward was doubled respectively to the previous one obtained. Session was terminated after 2h or when animals failed to lever press for three consecutive minutes, allowing to measure the session duration. Mice were tested multiple times in PRx2 with RR20 sessions intercalated between each PRx2 task to prevent operant responding extinction. For each animal, sessions of PRx2 were averaged, excluding the first one that was consistently an outlier.

#### PRx2 under amphetamine or CNO

Amphetamine (1 mg/kg) or clozapine-N-oxide (CNO, 2 mg/kg) were administered intraperitoneally 10 min or 45 min before the beginning of the PRx2, respectively. In the first PRx2 task, half of the animals received saline (NaCl 0.9%) and the other half received amphetamine or CNO. A second PRx2 task was performed 1 week later, and the order of saline-amphetamine/CNO administration was counterbalanced within groups.

### Fiber photometry *in vivo* recordings

#### Setup configuration

RWD (R820 Fiber Photometry) fiber photometry setup was used, coupled to an operant cage with the same configurations as those described above. A 470nm green-fluorescent protein (GFP) excitatory LED light stimulation coupled to a 410nm isosbestic stimulation was delivered through a fiber-optic patch cord (O.D. 1.25mm, Ceramic Ferrule, 400µm core diameter, 0.5 NA, R-FC-L-N5-400-L1, RWD). Fluctuations in fluorescence, reflecting variations of DA transmission, were computed and digitalized on the Multichannel Fiber Photometry Software (RWD, v2.0.0.33169).

#### Operant protocol

Mice were first trained in Pavlovian conditioning for 15-20 days. During the 15 min session, the emission of a light and sound stimulus (Conditioned Stimulus, CS) predicted the delivery of a drop of diluted sweet milk (Unconditioned Stimulus, US) (15 µl doses, diluted in water at 17.5%, Nestlé). The CS was on for 10 seconds, and the US occurred 3 seconds after the CS onset. Once the trough was empty and the CS was off, a new trial began. The inter-trial interval (ITI) lasted 30 seconds. Animals were then trained in Fixed Ratio 1 (FR1), during which both levers were out but only one was reinforced. When the RL was pressed, both levers retracted, and the CS started with the same pattern as Pavlovian training. Levers were out again at the beginning of the new trial, after an ITI of 10 seconds. After FR1 training, animals were trained in FR5 and FR10. All fixed-ratio sessions lasted 20 min.

#### Analysis

Fiber photometry recordings of DA transients in the NAc during Pavlovian and operant tasks were analyzed using a MATLAB custom script. The isosbestic signal was fitted to the 470 nm signal, and ΔF/F was calculated as: 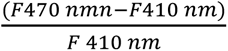, fitted for each time point. The z-score around each behavioral event was then calculated using a baseline of X sec (from - x sec to - x sec before the event) as:

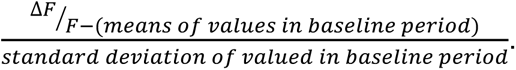

### Sample collection

On one side, mice blood was taken from the submandibular vein, and immediately after the animals were euthanized by cervical dislocation. All blood samples were centrifuged (15 min, 3500 rpm, 4°C), and plasma was collected and stored at - 80°C. The liver was collected and immediately stored at - 80°C. The entire brain was also collected and immediately frozen in isopentane and stored at - 80°C. Samples of dorsal striatum (DS), NAc and VTA were obtained from 1.5- and 1-mm punches of brain, respectively. To do so, 200 µm coronal brain sections were performed using a cryostat tissue slicer (Leica). According to the mouse brain atlas (40), punches of the DS were taken between 0.86 and 0.14 mm from the bregma, between 1.70 and 0.98 mm for the NAc and between - 2.92 and - 3.88 mm for the VTA. Punches were immediately stored at - 80°C.

On the other side, mice received an injection of buprenorphine (BUPRECARE®, 0.1 mg/kg) for analgesia, 30 min before the procedure. Mice were then injected with a lethal dose of pentobarbital (Exagon, 300 mg/kg) and intracardially perfused with cold Phosphate Buffer (PBS 1X) to remove blood, then by a cold 4% paraformaldehyde (PFA) solution (PFA in PBS, pH=7.4) in order to fix brain tissues. Brains were then coronally sectioned at 40 µm on a vibratome (Leica), collected and conserved in a cryoprotective solution (20% glycerol, 30% ethylene glycol, 50% PBS 1X, pH=7.4) at - 20°C until processing.

### Retinol and retinyl esters assays

The amount of retinol and retinyl esters were measured in plasma and liver as described in (41). Briefly, the molecules of interest were first extracted in an organic phase. The organic phase was left to evaporate under nitrogen until obtaining a dried extract. All dried extracts were dissolved in 200 µL HPLC mobile phase. A volume of 50-180 µL was used for HPLC analysis. All compounds were separated using a 250 x 4.6 mm RP C18, 5 µm Zorbax Eclipse XDB-C18 (Agilent Technologies, Santa Clara, CA, USA) preceded by a guard column, maintained at a temperature of 35°C. The mobile phase consisted of acetonitrile-dichloromethane-methanol (70:20:10; vol:vol:vol), using an isocratic elution and a flow rate of 1.8 mL/min. The HPLC system comprised a separation module and photodiode array detector (Shimadzu, Marne-la-Vallée, France). Compounds were detected at their maximum absorption wavelengths, i.e., 325 nm for retinol and retinyl esters. Quantifications were performed using Chromeleon software (version 6.8).

### Real-Time quantitative Polymerase Chain Reaction

The expression of the different genes of interest from brain punches was conducted as described in (42). Briefly, total RNA was extracted from brain punches using TRIzol® reagent (Invitrogen, Fisher Scientific, France). The purity and quantity of RNAs obtained were measure by using a Nanodrop One Spectrophotometer (Life technologies, France), and their quality was assessed using the RNA 600 Nano LabChip kit in combination with the 2100 Bioanalyzer (Agilent Technologies, France). Next, 500 ng of RNA were reverse-transcribed intro complementary DNA using the enzyme ImProm-II reverse transcriptase (A3802, Promega, France) in a final volume of 20 µL. Then, the RT-qPCR was performed in duplicate for each sample, with 1:50 cDNA diluted in a final volume of 10 µL, deposited in 384-well plated, and using the LightCycler 480 II system (Roche Diagnostics, Meylan, France). Gene amplication was performed using a SYBR Green I Master kit (RR240L, TAKARA BIO Europe, France) and appropriate primers. Data were analyzed using the comparative threshold cycle method performed with GenEx version 7 software (MultiD Analyses AB, Göteborg, Sweden). Results were normalized with 2 house-keeping genes, B2-microglobulin (B2M) and glyceraldehyde 3-phosphate dehydrogenase (GAPDH) and expressed as percentage relative expression of mRNA, with the mean of the control animals equal to 100%. The forward and reverse primer sequences used to amplify genes of interest are summarized in Supplementary Table 1.

### Western blot

Proteins were extracted from tissues using Tissue Lyser (Qiagen) in extraction buffer containing 50 mM Tris, 0.5 M urea, 2% SDS (Sodium Dodecyl Sulfate) and protease and phosphatase inhibitors (Fisher Scientific). After sonication and centrifugation, the total protein concentration of the supernatant was assessed using the BC Assay protein quantification kit (Interchim) following the manufacturer’s instructions. An equivalent amount of protein (7,5 µg/well) and a molecular weight marker (Thermo Scientific, PageRuler^TM^) were separated by electrophoresis in a polyacrylamide gel (10%) before being transferred to a nitrocellulose membrane (Amersham Protan Premium 0.2 µm). Membranes were saturated in a solution of Intercept® LI-COR for 1h. After a 5 min wash in TBS-Tween (20 0.1%, Sigma), membranes are then incubated overnight at 4°C with the primary antibodies. After several washes in TBS 1X (Tris-Buffered Saline), the membranes were incubated with the secondary antibodies for 1h at room temperature protected from the light. Antibodies were diluted in 1% (Régilait) milk in TBS 1X. Fluorescence was detected using the Odyssey® CLx LI-COR imaging system. Signal intensity was quantified with the Image Studio® software and normalized to the housekeeping protein (α-tubulin, 50 kDa) expression for each sample. Results are expressed in % relative to control animals. Antibodies used are summarized in Supplementary Table 1.

### Statistical analysis

All data are displayed as means ± SEM with single data points plotted. Data were analyzed with GraphPad Prism 10. Statistical significance was *p<0.05, **p<0.01, ***p<0.001, ****p<0.0001. Detailed statistical analysis and results are reported in Supplementary Table 3.

## RESULTS

### Developmental VAD enhances motivational performance in male offspring

We first assessed the effect of VAD from gestation to adulthood (Figure 1A) on motivational processes, using operant conditioning paradigms (Figure 1B). Performances evaluated by the number of lever presses on the RL to obtain palatable food rewards across increasing random ratio requirements were similar between CD and VAD mice for both sexes (Figure 1C). To evaluate motivational performance, we used the PRx2 task which assessed the animal’s motivation to maintain effort in the face of an exponentially increasing demand (Figure 1D). Developmental VAD induced a significant increase in total lever-pressing and breakpoint (the last progressive ratio attained) only in male offspring, while females were not impacted (Figure 1E-F). Importantly, neither preference for palatable sucrose solution – as a proxy for hedonic reactivity - nor spontaneous locomotion were altered in VAD mice (Supplementary Figure 1A-B). Moreover, the ability to adapt behavior to a change in outcome value was spared, supporting that goal-directed behavior was preserved (Supplementary Figure 1C). Considering the relationship between motivation and impulsivity (43–47) and the tendency to an increased lever-pressing on the unreinforced lever (UL) in the PRx2 task in VAD male mice (Supplementary Figure 1D), we assessed choice impulsivity using the delay discounting task. VAD male switched faster onto the lever associated to the low but immediate reward, demonstrating elevated impulsivity. VAD females were not affected in this paradigm (Supplementary Figure 1E-F).

**Figure 1.**
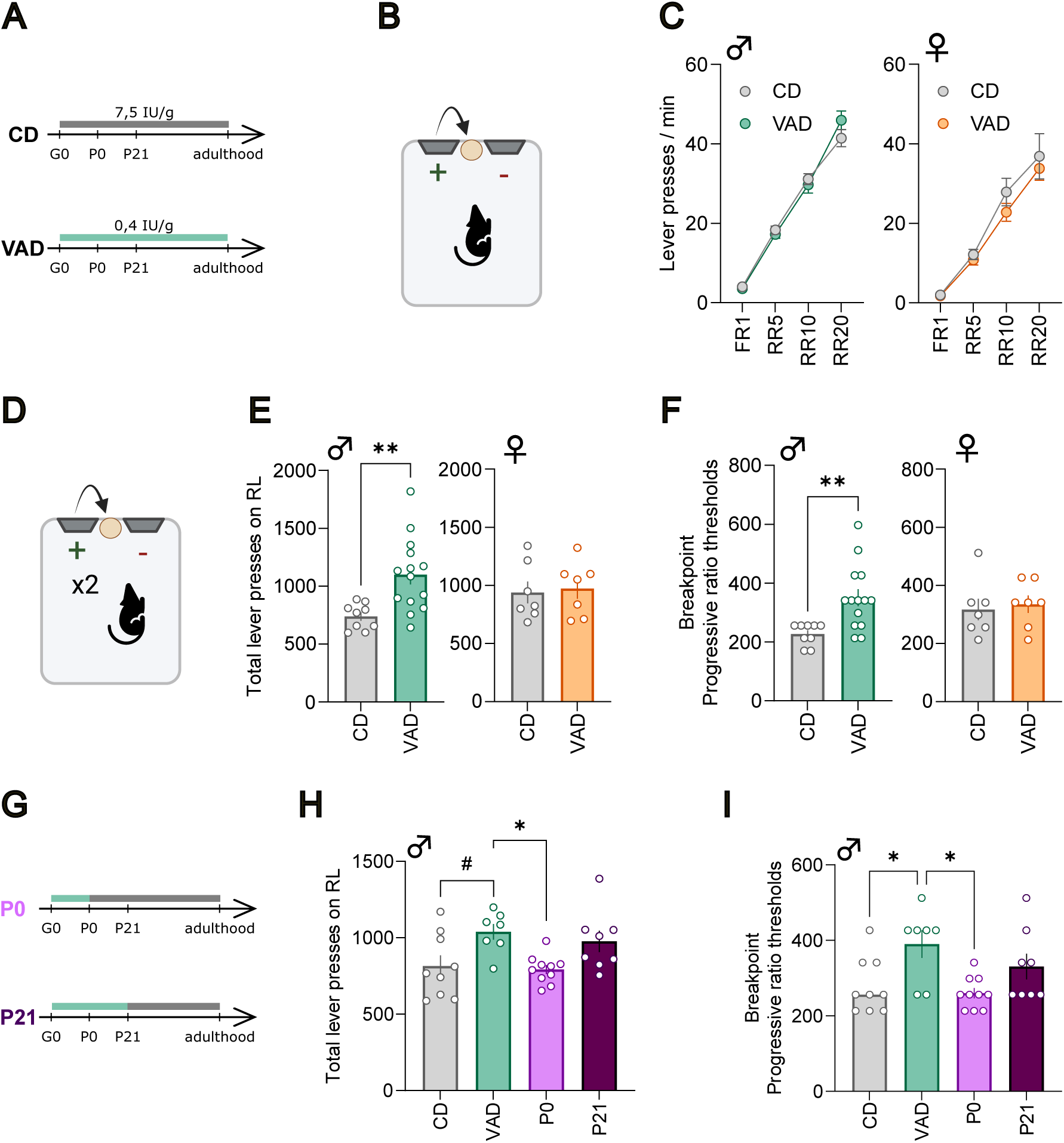
Developmental VAD enhances motivational performance in male offspring. **(A)** Schematic representation of the two experimental groups. Control diet (CD) contains 7.5 IU of retinol/g while vitamin A deficient diet (VAD) contains 0.4 IU of retinol/g. Diet exposure started 4 weeks before mating in mothers and was maintained throughout life in offspring. **(B)** Schematic illustration of the operant conditioning paradigm. Mice must press on a reinforced lever (RL) to obtain a palatable food reward. Pressing the unreinforced lever (UL) has no consequences. **(C)** Lever pressing rate (lever presses/min) as a function of operant ratio in male (two-way ANOVA: diet *p=0.7418*, ratio \*\*\*\**p<0.0001* and interaction *p=0.2385*; CD: n=9, VAD: n=15) and female (two-way ANOVA: diet *p=0.2234*, ratio \*\*\*\**p<0.0001* and interaction *p=0.2385*; CD: n=7, VAD: n=7) offspring. **(D)** Paradigm for the progressive ratio task, where the number of lever presses required to obtain a reward is doubled respectively to the previous one obtained. **(E)** Total lever presses on the RL (unpaired t-test: 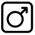 \*\**p=0.0039*; 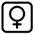 *p=0.7958*) and **(F)** breakpoint (i.e., last progressive ratio threshold reached by the mice) (Mann-Whitney test: 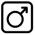 \*\**p=0.0018*; unpaired t-test: 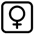 *p=0.7087*) in the progressive ratio (x2) in male (CD: n=9, VAD: n=15) and female (CD: n=7, VAD: n=7) offspring. **(G)** Schematic representation of the two experimental supplementation groups at either P0 or P21. **(H)** Total lever presses on the RL (one-way ANOVA: \**p=0.0107*; Tukey’s multiple comparisons: CD vs VAD: #*p=0.0523*; VAD vs P0: \**p=0.0238*) in male (CD: n=9, VAD n=7, P0 n=10, P21 n=8). **(I)** Breakpoint (one-way ANOVA: \**p=0.0102*; Tukey’s multiple comparisons: CD vs VAD: \**p=0.0399*; VAD vs P0: \**p=0.0101*) in male (CD: n=9, VAD n=7, P0 n=10, P21 n=8). ANOVA: analysis of variance, CD: control diet, VAD: vitamin A deficient, IU: international unit, FR: fixed ratio, RR: random ratio, RL: reinforced lever, UL: unreinforced lever (Table S3 for detailed statistical analysis and results).

Importantly, we found that normalization of vitamin A levels at birth (P0), but not at weaning (P21) prevents hypermotivation at adulthood in VAD male mice (Figure G-I), supporting that altered reward processing at adulthood originates from developmental effects.

### Developmental VAD impacts peripheral and central RA signaling together with the expression of markers of DA transmission in the mesolimbic pathway

The behavioral alterations evidenced only in VAD males prompted us to interrogate potential sex-dependent differences in sensitivity to VAD, focusing on the mesolimbic pathway considering its established role in reward processing (48,49).

Developmental VAD did not modify body weight gain (data not shown) but reduced circulating plasma retinol levels in both male and female offspring (Figure 2A). Of note, plasma retinol concentrations were lower in females than in males in the CD group as previously reported (41). Similarly, free retinol and retinyl esters were strongly decreased by VAD in the liver, the storage organ of vitamin A, in both males and females (Figure 2B-C). These data suggest that VAD from gestation to adulthood similarly impacts peripheral vitamin A metabolism in both males and females.

**Figure 2.**
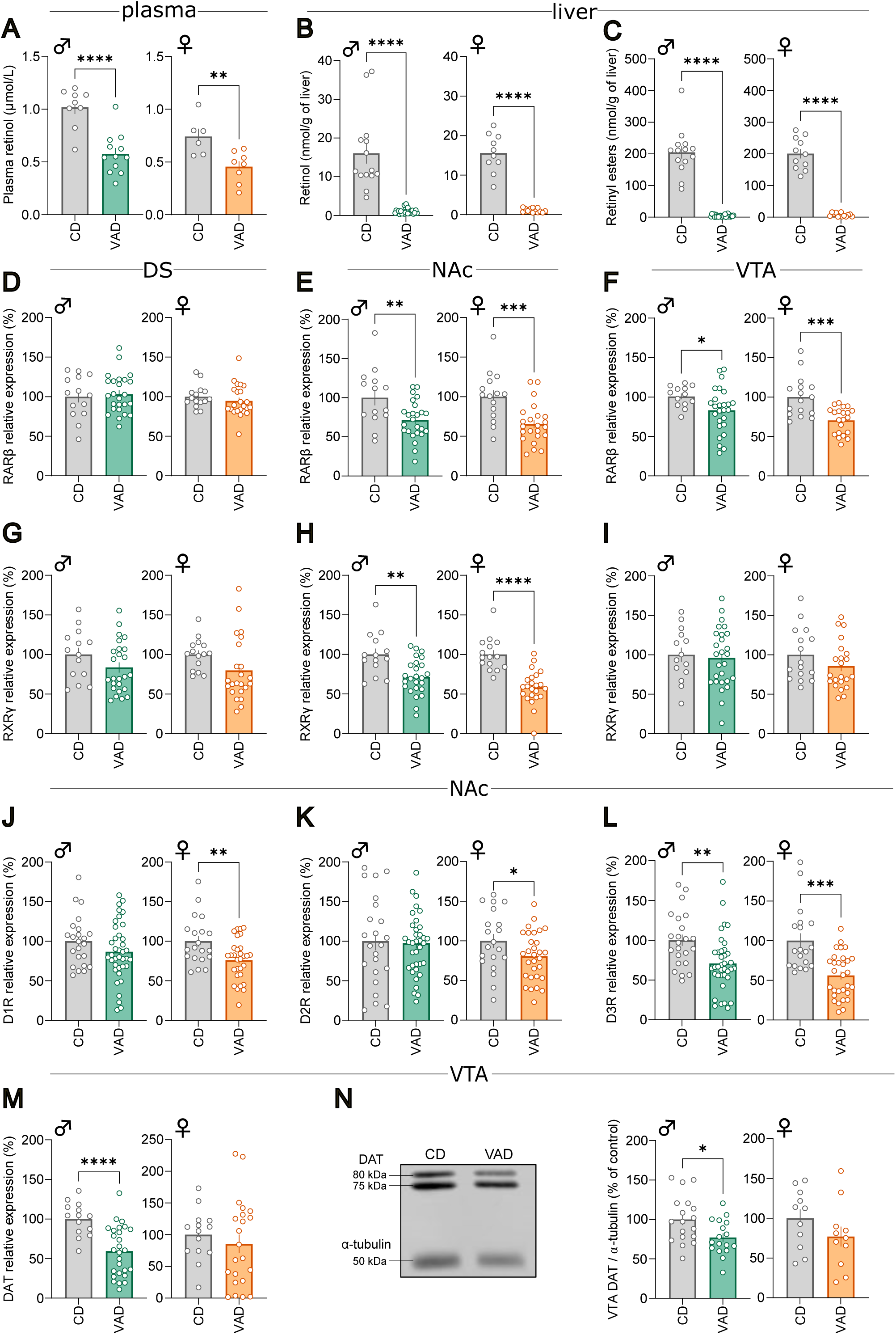
Developmental VAD impacts peripheral and central RA together with the expression of markers of DA transmission in the mesolimbic pathway. **(A)** Circulating plasma retinol (µmol/L) in male (unpaired t-test: \*\*\*\**p<0.0001*; CD: n=8, VAD: n=12) and female (unpaired t-test: \*\**p=0.0069*; CD: n=6, VAD: n=8) offspring. **(B)** Hepatic retinol (nmol/g of liver) in male (Mann-Whitney test: \*\*\*\**p<0.0001*; CD: n=14, VAD: n=25) and female (Mann-Whitney test: \*\*\*\**p<0.0001*; CD: n=11, VAD: n=13) offspring. **(C)** Hepatic retinyl esters (nmol/g of liver) in male (Mann-Whitney test: \*\*\*\**p<0.0001*; CD: n=14, VAD: n=25) and female (Mann-Whitney test: \*\*\*\**p<0.0001*; CD: n=11, VAD: n=13) offspring. **(D)** mRNA relative expression of genes coding for RARβ (unpaired t-test: 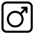 *p=0.7017*; Mann-Whitney test: 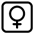 *p=0.2043*) and **(G)** RXRγ (unpaired t-test: 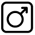 *p=0.1329*; Mann-Whitney test: 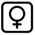 *p=0.0802*) in the dorsal striatum (DS) in male (CD: n=15, VAD: n=23) and female (CD: n=14, VAD: n=25) offspring. **(E)** mRNA relative expression of genes coding for RARβ (unpaired t-test: 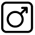 \*\**p=0.0058*; 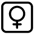 \*\*\**p=0.0006*) and **(H)** RXRγ (unpaired t-test: 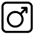 \*\**p=0.0012*; 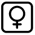 \*\*\*\**p<0.0001*) in the nucleus accumbens (NAc) in male (CD: n=15, VAD: n=23) and female (CD: n=14, VAD: n=25) offspring. **(F)** mRNA relative expression of genes coding for RARβ (unpaired t-test: 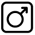 \**p=0.0426*; Mann-Whitney test: \*\*\**p=0.0009*) and **(I)** RXRγ (unpaired t-test: 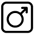 *p=0.7396*; 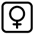 *p=0.1808*) in the ventral tegmental area (VTA) in male (CD: n=14, VAD: n=27) and female (CD: n=16, VAD: n=22) offspring. **(J)** mRNA relative expression of genes coding for D1R (unpaired t-test: 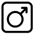 *p=0.1501*; 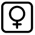 \*\**p=0.0045*), **(K)** D2R (unpaired t-test: 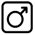 *p=0.8520*; 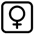 *p=0.0533*) and **(L)** D3R (Mann-Whitney test: 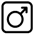 \*\**p=0.0020*; 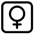 \*\*\*\**p=0.0003*) in the NAc in male (CD: n=23, VAD: n=39) and female (CD: n=20, VAD: n=30) offspring. **(M)** mRNA relative expression of the gene coding for DAT (unpaired t-test: 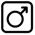 \*\*\*\**p<0.0001*; 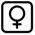 *p=0.4782*) in VTA in male (CD: n=14, VAD: n=27) and female (CD: n=15, VAD: n=22) offspring. **(N)** Representative western blot for DAT in male (left) and quantification of protein expression levels of DAT (right) (unpaired t-test: 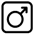 \**p=0.0154*; 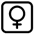 *p=0.1865*) in the VTA in male (CD: n=18, VAD: n=17) and female (CD: n=11, VAD: n=12) offspring. CD: control diet, VAD: vitamin A deficient, RAR: retinoic acid receptor, RXR: retinoid x receptor, DR: dopamine receptor, DAT: dopamine transporter, DS: dorsal striatum, NAc: nucleus accumbens, VTA: ventral tegmental area. (Table S2 for detailed statistical analysis and results). (Table S3 for detailed statistical analysis and results).

We then assessed mRNA expression of retinoid receptors (RAR, RXR) – as an indicator for brain RA signaling, since their expression are directly controlled by RA levels (31). We found no change in the DS, while expression of RARβ and RXRγ were decreased in the NAc, as well as in the VTA (for RARβ only) in both sexes (Figure 2D-I). Overall, these findings suggest that the mesolimbic pathway is particularly sensitive to VAD, in a similar way in males and females.

We then assessed the functional integrity of mesolimbic DA transmission by quantifying mRNA and protein expression of key markers. While female VAD mice displayed significant decreased mRNA expression of dopamine receptors (DR) D1R, D2R and D3R in the NAc, only D3R was downregulated in male mice (Figure 2J-L), with no change in expression in the DS for both sexes (Supplementary Figure 2A-C). When analyzing presynaptic markers of DA transmission in the VTA, there was no change in expression of genes coding for proteins involved in DA synthesis (tyrosine hydroxylase (TH), dopamine decarboxylase (DDC)), degradation (monoamine oxidase A (MAOA)) or vesicular storage (vesicular monoamine transporter 2 (VMAT2)) (Supplementary Figure 2D-G). However, we found a significant decrease in mRNA (Figure 2M) and protein levels (Figure 2N) of the DA transporter (DAT) in the VTA of VAD males, but not females. Interestingly, RARβ expression positively correlated with DAT expression in the VTA in males (Supplementary Figure 2H). Importantly, reduced expression in VAD male mice did not result from a decrease in the number of TH-positive DA neurons in the VTA (Figure S2I-J) nor in DA fiber density in the NAc (Supplementary Figure 2K-L), as revealed by TH immunofluorescent staining.

### Developmental VAD potentiates phasic DA dynamics in the NAc of male offspring in relation with effort expenditure

Decreased expression of DAT selectively in male VAD mice prompted us to investigate the integrity of mesolimbic DA dynamics. Indeed, hyperdopaminergia due to DAT loss-of-function in DA neurons is associated with enhanced operant responding as well as increased impulsivity (50–56), which resembles the behavioral phenotype of VAD male mice. We therefore evaluated DA dynamics using the DA sensor dLight coupled with fiber photometry (Figure 3A), in the NAc core of male VAD mice (Figure 3B), in relation to discrete behavioral epochs associated with reward prediction and consumption, as well as effort evaluation and engagement. Mice were first trained in a Pavlovian conditioning task where a cue (CS, i.e., a sound and light on for 10 seconds) predicts the delivery of a reward (US), 3 seconds after the beginning of the CS (Figure 3C). There was no difference in DA dynamics aligned to the CS or the US (Figure 3D-E), suggesting that DA encoding of reward prediction (CS) and reward value (US) were spared in VAD animals. Mice were then trained in a FR1 task, in which one lever press on the RL triggered the CS-US schedule, followed by FR5 (5 lever presses) and FR10 (10 lever presses). No significant changes were observed in the response to the US (Supplementary Figure 3A-B). However, DA responses at the CS were increased across FRs in VAD mice (Figure 3F-G). Surprisingly, while virtually absent in control animals, there was a significant DA response when lever pressing on the UL in VAD mice (Supplementary Figure 3C-D). Moreover, when looking at DA dynamics at the first lever press of bouts – i.e. when animals engage in effort –, we found enhanced responses in VAD male mice (Figure 3H-I). Regarding mice behavior, the number of licks, reward, RL and UL presses were not different between CD and VAD in Pavlovian (Supplementary Figure 3E) and FRs (Supplementary Figure 3F).

**Figure 3.**
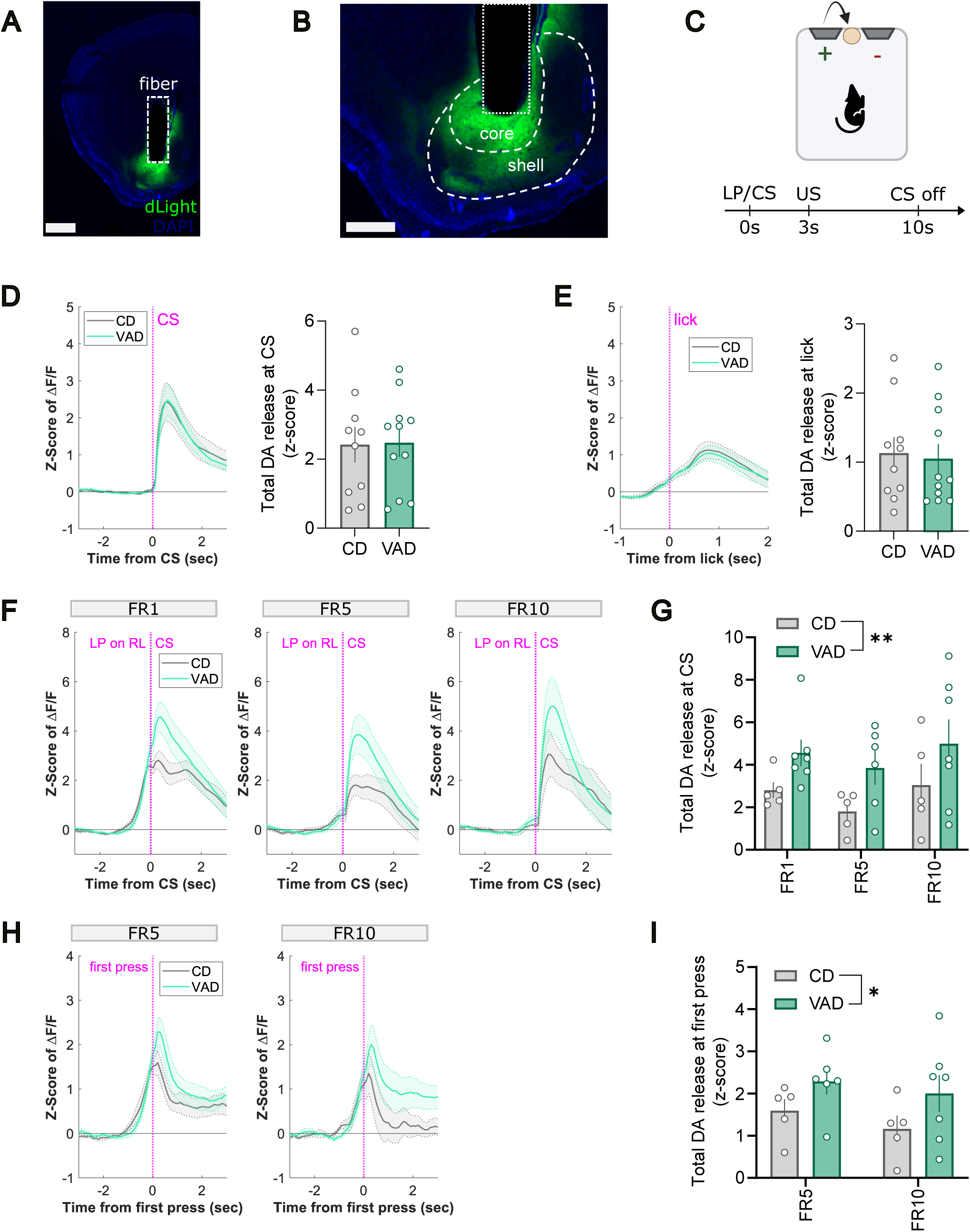
Developmental VAD potentiates phasic dopamine dynamics in the nucleus accumbens of male offspring in relation with effort expenditure. **(A)** Unilateral injection of a viral vector allowing dLight expression in the nucleus accumbens (NAc) of WT mice with a representative example of viral infection (green fluorescent protein (GFP) staining) and optic fiber implantation, and nuclear staining with DAPI (scale bar=800µm). **(B)** Zoom of the viral spread in the NAc (scale bar=250µm). **(C)** Timeline of a trial. Each trial started with the onset of the conditioned stimulus (CS) after a reinforced lever (RL) press (for fixed ratio trials) or not (for Pavlovian conditioning trials), followed by the milk delivery (US) after 3 seconds. After another 7 seconds (10 seconds in total), the cue was turned off. **(D)** Dopamine (DA) dynamic (z-score of ΔF/F) during Pavlovian conditioning aligned to the CS (left) and quantification of the peak amplitude (z-score, 0 to 2 seconds) (unpaired t-test: *p=0.9317*) in males (CD: n=10; VAD: n=11). **(E)** DA dynamic during Pavlovian conditioning around licking onset (left) and quantification of the peak amplitude (z-score, 0 to 2 seconds) (Mann-Whitney test: *p=0.7564*) in males (CD: n=10; VAD: n=11). **(F)** DA dynamic during fixed ratio (FR1, FR5 and FR10) trials aligned onto the RL press that triggers the CS in males (CD: n=5; VAD: n=7). **(G)** Quantification of DA dynamics (peak amplitude, z-score) (two-way ANOVA: diet \**p=0.0090*, ratio *p=0.3617* and interaction *p=0.9846*) at the CS (0 to 2 seconds) for the different fixed ratio trials in males (CD: n=5; VAD: n=7) relative to **(F)**. **(H)** DA dynamic during fixed ratio (FR5, FR10) trials aligned onto the first lever press of a bout in males (CD: n=5, VAD: n=7) **(I)** and quantification of the peak amplitude (0 to 2 sec) (two-way ANOVA: *diet *p=0.0499*, ratio *p=0.3429* and interaction *p=0.8516*). ANOVA: analysis of variance, CD: control diet, VAD: vitamin A deficient diet, FR: fixed ratio, LP: lever press, CS: conditioned stimulus, US: unconditioned stimulus, ITI: inter-trial interval, NAc: nucleus accumbens, DA: dopamine, RL: reinforced lever. (Table S3 for detailed statistical analysis and results).

Altogether, these results reveal enhanced mesolimbic DA dynamics in VAD male mice, suggesting a hyperdopaminergic phenotype.

### Inhibition of the mesolimbic pathway normalizes motivational performance in VAD males

Decrease in DAT expression classically dampens behavioral responding to the dopaminomimetic amphetamine (56–58). To link decreased DAT expression to increased operant responding in VAD male mice, we challenged animals with amphetamine and evaluated the impact on performance in PRx2 tasks. In accordance with its pro-motivational effect, amphetamine potentiated lever pressing in CD animals. By contrast, this effect was blunted in VAD male mice (Figure 4A, Supplementaru Figure 4A), supporting dampened DAT activity.

**Figure 4.**
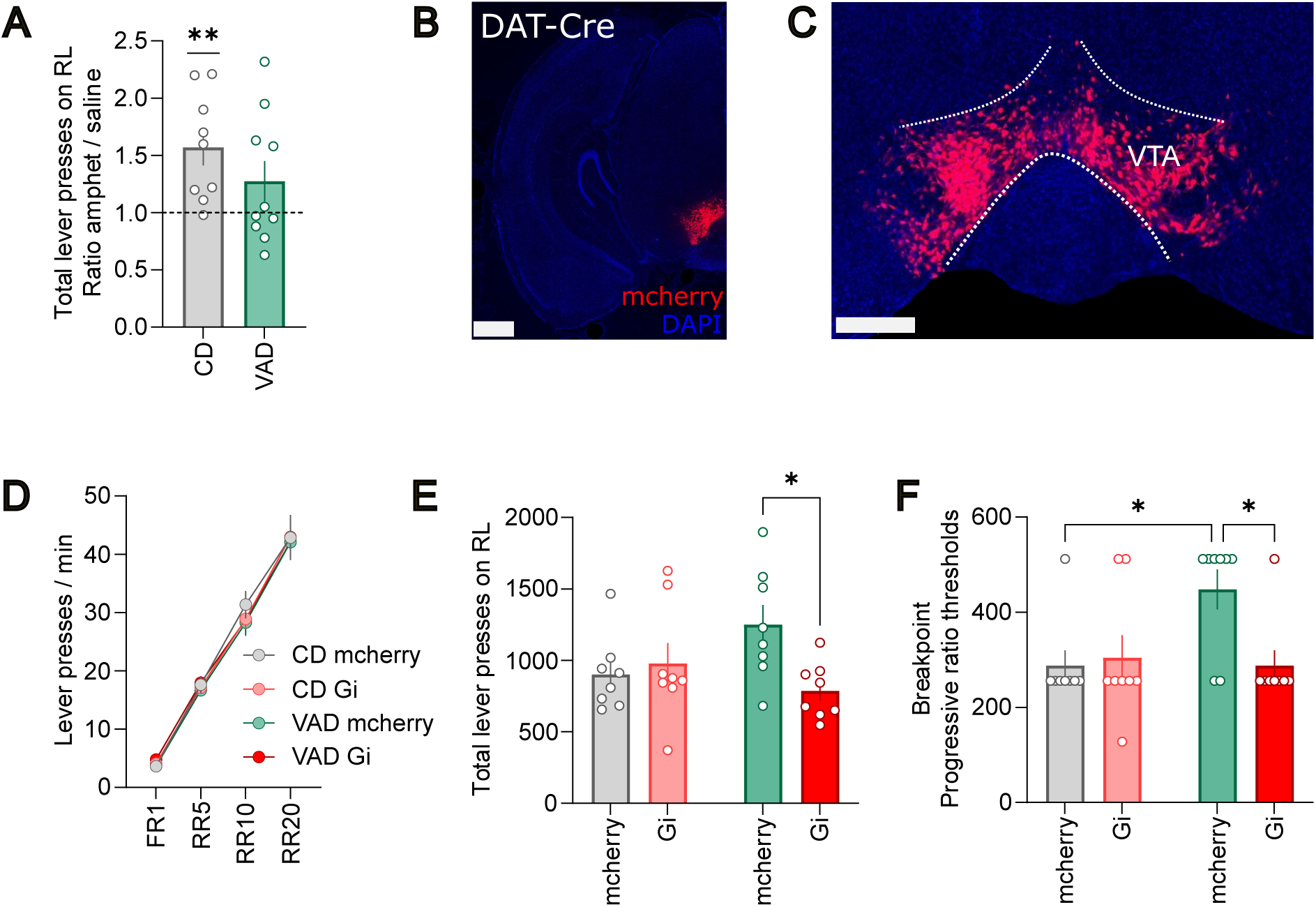
Inhibition of the mesolimbic pathway normalizes motivational performance in VAD males. **(A)** Ratio of total lever presses on the reinforced lever (RL) during the progressive ratio (PRx2) task under amphetamine (1 mg/kg, injected 10 min before the beginning of the PRx2 task) over saline, in male (CD: n=9, VAD: n=10) offspring (one-sample Wilcoxon test: CD \*\**p=0.0078*, VAD *p=0.3887*). **(B)** Bilateral injection of a viral vector allowing HM4Di expression in the ventral tegmental area (VTA) of DAT-Cre mice with a representative example of viral infection (mcherry staining) and nuclear staining with DAPI (scale bar=800µm). **(C)** Zoom of the viral spread in the VTA (scale bar=400µm). **(D)** Lever pressing rate (lever presses/min) on the RL during the instrumental learning phase (three-way ANOVA: diet *p=0.5835*, virus *p=0.9272*, ratio \*\*\*\**p<0.0001*, diet x virus *p=0.4842*, diet x ratio *p=0.8952*, ratio x virus *p=0.9270* and diet x virus x ratio *p=0.9525*) in males (mcherry CD: n=8, mcherry VAD: n=8, Gi CD: n=8, Gi VAD: n=8) without clozapine-n-oxide (CNO) injection. **(E)** Total lever presses on the RL (two-way ANOVA: diet *p=0.5041*, virus *p=0.1040* and interaction \**p=0.0269*; Tukey’s multiple comparisons: mcherry CD vs mcherry VAD: #*p=0.1684*; mcherry VAD vs Gi VAD: \**p=0.0392*) and **(F)** breakpoint (two-way ANOVA: diet *p=0.0760*, virus *p=0.0760* and interaction \**p=0.0323*; Tukey’s multiple comparisons: mcherry CD vs mcherry VAD: \**p=0.0345*; mcherry VAD vs Gi VAD: \**p=0.0345*) in the PRx2 task under CNO, in male offspring (mcherry CD: n=8, mcherry VAD: n=8, Gi CD: n=8, Gi VAD: n=8). ANOVA: analysis of variance, CD: control diet, VAD: vitamin A deficient, CNO: clozapine-n-oxide, FR: fixed ratio, RR: random ratio, VTA: ventral tegmental area, RL: reinforced lever. (Table S3 for detailed statistical analysis and results).

We finally tested whether the hyperdopaminergic phenotype in VAD males could be directly responsible for enhanced operant responding by assessing whether chemogenetic inhibition of VTA DA neurons could normalize behavioral performances in VAD males. Thus, AAV carrying a Cre-dependent inhibitory DREADD hM4Di or control mCherry fluorescent protein were injected in the VTA of DAT-Cre CD and VAD male mice (Figure 4B-C, Supplementary Figure 4B). All groups performed similarly in operant learning and in RR schedules in the absence of chemogenetic inhibition (Figure 4D). Activation of the DREADD by administration of the DREADD exogenous ligand, CNO (2 mg/kg) normalized lever pressing and breakpoint of VAD male mice (Figure 4E-F). Because of the lack of effects of our chemogenetic approach in CD mice, we used another motivational task, the concurrent lever pressing/free-feeding task, that assess the willingness to exert effort for a preferred reward. This paradigm demonstrated an overall effect of the viral approach, confirming the effectiveness of our chemogenetic inhibition (Supplementary Figure 4B).

Overall, these results demonstrated a causal link between hyperactivity of the mesolimbic DA transmission and enhanced operant responding in VAD male offspring.

## DISCUSSION

Our findings provide strong evidence for sex-specific vulnerability of the mesolimbic DA transmission to developmental VAD in mice, leading to altered reward processing at adulthood. We demonstrate that increased operant responding in VAD male mice results from hyperdopaminergia, which relates, at least in part, from decreased DAT expression in midbrain DA neurons. Indeed, DAT loss-of-function has been largely shown to enhance motivation and impulsivity (50–56) which resembles the behavioral phenotype of VAD male mice. In accordance with DAT loss-of-function leading to hyperdopaminergia (59,60), our fiber photometry data based on expression of the DA sensor in the NAc show an increased DA dynamic specifically in effort-based tasks. Altogether, these findings further support hyperdopaminergia as a main neurobiological mechanism for hypermotivation in VAD male mice, which is in accordance with the ability of chemogenetic inhibition of VTA DA neurons to normalize operant responding.

While the direct link remains to be established, we hypothesize that DAT downregulation is a major factor by which VAD induces hyperdopaminergia. Indeed, RA hyposignaling correlates with decreased DAT expression, and amphetamine injection, known to increase extracellular DA levels, particularly in the NAc (61), fails to exert its pro-motivational effects. While this could be due to a ceiling effect on DA levels, it is more likely that the efficacy of amphetamine to act as an inhibitor of DAT is blunted due to decreased DAT expression. The mechanisms by which VAD decreases DAT expression are unknown, however it has been shown that high vitamin A intake during pregnancy leads to increased hippocampal DAT expression in offspring through epigenetic changes (62). We cannot rule out that other mechanisms might be at play in mediating altered hyperdopaminergia-related reward processing in VAD male mice. Notably, we found that DA receptors expression – that is modulated by RA (63) - is decreased in the NAc of VAD animals, in accordance with the effect of retinoid receptors loss-of-function (33,35,36,64). Expression of DA receptors has been well established to influence both motivation (65–68) and impulsivity (69–72). However, VAD females show downregulation of all DA receptors in the NAc whereas they do not display alterations in motivation. Moreover, D3R expression is decreased in VAD males, which is unlikely to mediate enhanced motivational performance since genetic invalidation of D3R in the NAc has rather been associated with motivational deficits (73).

RA signaling has been largely involved in striatal neurogenesis (64) and differentiation of medium spiny neurons (26,27). Importantly, our data suggest that the effects of developmental VAD on reward processing in male offspring take place during the perinatal period since vitamin A supplementation from birth can prevent behavioral alterations in VAD male offspring, but not supplementation from weaning. Moreover, our data show higher vulnerability of the NAc as compared to the DS to RA signaling, which is consistent with transcriptomic analysis of RARβ mutant mice that revealed a dorso-ventral gradient with drastic alterations in gene expression in the NAc compared to the DS (36). RA signaling has been shown to be also critical for differentiation of DA neurons (25,74) even if its role in the development and functionality of DA transmission *per se* remains poorly explored. While we did not observe gross alteration in the density of DA neurons or projections, we found decreased DAT expression in the VTA in VAD male mice. However, it remains to be clarified whether this decreased DAT expression during development is the primary cause or a consequence of DA dysfunction. Lack of evidence shows the involvement of DAT for the initial development of DA neurons *per se*. However, studies show that DAT is present in the developing brain as early as the late gestational period and then declining with age, suggesting a role in neuronal and synaptic maturation (75,76). Moreover, in the developing neurons, DAT is dynamically trafficked to the plasma membrane of developing dopaminergic axons, particularly concentrating in growth cones and filopodia—structures closely associated with synaptogenesis (77). Further analyses will be necessary to assess the ultrastructure of DA synapses in VAD mice, and whether this could be at the core of hyperdopaminergia and related behavioral alterations.

The question remains concerning the resistance of female offspring to VAD, at least regarding reward processes and DA transmission. However, available evidence of the effects of VAD on brain function focuses on male animals and does not directly address sex-specific differences. Nonetheless such sex difference is particularly relevant regarding the implication of RA signaling in the etiology of neurodevelopmental disorders for which elevated striatal DA is a main neurobiological feature, such as schizophrenia or bipolar disorders (78–80). Indeed, schizophrenia has been shown to be prevalent in boys, with an earlier age of onset, greater symptom severity, and a poorer response to treatment (81–83). Attention deficit hyperactivity disorder is also more common in boys and has different clinical manifestations in men and women (81,84,85). This had led to the “estrogen hypothesis” which postulates that estrogens provide neuroprotective effects regarding the onset, progression and symptom severity of psychiatric disorders (83,86–88). The effect of ovarian hormones on the regulation of DA transmission by developmental RA signaling remains to be clarified.

Altogether, we demonstrated for the first time that blunted RA signaling induced by nutritional VAD during brain development induces sex-specific motivational processing alterations, through a dysregulation of the mesolimbic DA transmission in male adult offspring. Thus, our data support that RA signaling appears as promising target and relevant transdiagnostic biomarker to better predict and improve treatments for some neurodevelopmental psychiatric disorders. Further preclinical studies will be needed to better understand mechanisms through which RA hyposignaling hijacks the development of DA transmission, hence leading to incentive processing alterations.

## ACKNOWLEDGMENTS AND DISCLOSURES

We thank the Bordeaux Imaging Center (a service unit of the CNRS-INSERM and Bordeaux University, member of the national infrastructure France BioImaging) support by ANR-10-INBS-04 for imaging, Céline Ducroix-Crépy (INRAE UMR 1286) for genotyping, the staff from the animal facility of INRAE UMR 1286 for animal care and the CIRCE (Behavioural Engineering Center) facility of the Bordeaux Neurocampus.

This study was supported by INRAE and University of Bordeaux, University of Bordeaux’s IdEx “Investments for the future” program/GPR BRAIN_2030 (PT), ANR “FrontoFat” (ANR-20-CE14-0020) (PT), ANR “StriaPOM” (ANR-21-CE14-0018) (PT), ANR “BRAINHEALTH” (ANR-23-CE14-0018) (PT), Institut de Recherche en Santé publique (IReSP) Aviesan APP-addictions 2019 (PT), Institut de Recherche en Santé publique (IReSP) APP-addictions 2023 (PT), Labex “BRAIN” (PT and RW), Region Nouvelle Aquitaine 2014-1R30301-00003023 (PT); Fondation pour la Recherche Médicale (FRM “Environnement et Santé”) (GF and PT), Equipe FRM EQU202403018022 (PT), ANSSD 2023 INRAE Département AlimH (KT and PB). PC, RW and LH were the recipient of a PhD fellowship from the French Ministry of Research and Higher Education and AP from the “Ecole Universitaire de Recherche” (EUR Neuro, Bordeaux Neurocampus). ID and LB were the recipient of a Master fellowship from APP-Bordeaux INP 2023 and 2024.

PC, KT and PT conceived the project, designed the study and wrote the manuscript. PC, SY, ID, LB and AS performed most of the experiments, analyzed and interpreted the data. SA, JCH and MFA provided help and training for experiments. LH, AP, FD, RW and JC developed behavioral tasks and fiberphotometry analyses. CS and PB performed the retinol and retinyl esters assays. PB, GF and CBB discussed the data and provide inputs to experiments and manuscripts. All authors read, edited and approved the final manuscripts.

The authors report no biomedical financial interests or potential conflicts of interest.

## REFERENCES

1. Whitton AE, Treadway MT, Pizzagalli DA. Reward processing dysfunction in major depression, bipolar disorder and schizophrenia. Current Opinion in Psychiatry. janv 2015;28(1):7-12.

2. Husain M, Roiser JP. Neuroscience of apathy and anhedonia: a transdiagnostic approach. Nat Rev Neurosci. août 2018;19(8):470-84.

3. Moeller FG, Barratt ES, Dougherty DM, Schmitz JM, Swann AC. Psychiatric Aspects of Impulsivity. AJP. 1 nov 2001;158(11):1783-93.

4. Kulacaoglu F, Kose S. Singing under the impulsiveness: impulsivity in psychiatric disorders. Psychiatry and Clinical Psychopharmacology. 3 avr 2018;28(2):205-10.

5. Dalley JW, Roiser JP. Dopamine, serotonin and impulsivity. Neuroscience. juill 2012;215:42-58.

6. Maden M. Retinoic acid in the development, regeneration and maintenance of the nervous system. Nat Rev Neurosci. oct 2007;8(10):755-65.

7. Goodman AB. Three independent lines of evidence suggest retinoids as causal to schizophrenia. Proc Natl Acad Sci USA. 23 juin 1998;95(13):7240-4.

8. Bao Y, Ibram G, Blaner WS, Quesenberry CP, Shen L, McKeague IW, et al. Low maternal retinol as a risk factor for schizophrenia in adult offspring. Schizophrenia Research. mai 2012;137(1-3):159-65.

9. Wan C, Shi Y, Zhao X, Tang W, Zhang M, Ji B, et al. Positive association between ALDH1A2 and schizophrenia in the Chinese population. Progress in Neuro-Psychopharmacology and Biological Psychiatry. nov 2009;33(8):1491-5.

10. Regen F, Cosma NC, Otto LR, Clemens V, Saksone L, Gellrich J, et al. Clozapine modulates retinoid homeostasis in human brain and normalizes serum retinoic acid deficit in patients with schizophrenia. Mol Psychiatry. sept 2021;26(9):5417-28.

11. Yang CD, Cheng ML, Liu W, Zeng DH. Association of serum retinoic acid with depression in patients with acute ischemic stroke. Aging (Albany NY). 10 févr 2020;12(3):2647-58.

12. Li HH, Yue XJ, Wang CX, Feng JY, Wang B, Jia FY. Serum Levels of Vitamin A and Vitamin D and Their Association With Symptoms in Children With Attention Deficit Hyperactivity Disorder. Front Psychiatry. 23 nov 2020;11:599958.

13. Wang CX, Wang B, Sun JJ, Xiao CY, Ma H, Jia FY, et al. Circulating retinol and 25(OH)D contents and their association with symptoms in children with chronic tic disorders. Eur Child Adolesc Psychiatry. avr 2024;33(4):1017-28.

14. Feng J, Chen J, Yan J, Jones IR, Craddock N, Cook EH, et al. Structural variants in the retinoid receptor genes in patients with schizophrenia and other psychiatric diseases. American J of Med Genetics Pt B. 5 févr 2005;133B(1):50-3.

15. Brzustowicz LM, Hodgkinson KA, Chow EWC, Honer WG, Bassett AS. Location of a Major Susceptibility Locus for Familial Schizophrenia on Chromosome 1q21-q22. Science. 28 avr 2000;288(5466):678-82.

16. Galter D, Buervenich S, Carmine A, Anvret M, Olson L. ALDH1 mRNA: presence in human dopamine neurons and decreases in substantia nigra in Parkinson’s disease and in the ventral tegmental area in schizophrenia. Neurobiology of Disease. déc 2003;14(3):637-47.

17. Wo\loszynowska-Fraser MU, Kouchmeshky A, McCaffery P. Vitamin A and Retinoic Acid in Cognition and Cognitive Disease. Annual Review of Nutrition. 2020;40(1):247-72.

18. Qi XR, Zhao J, Liu J, Fang H, Swaab DF, Zhou JN. Abnormal Retinoid and TrkB Signaling in the Prefrontal Cortex in Mood Disorders. Cerebral Cortex. janv 2015;25(1):75-83.

19. Reay WR, Atkins JR, Quidé Y, Carr VJ, Green MJ, Cairns MJ. Polygenic disruption of retinoid signalling in schizophrenia and a severe cognitive deficit subtype. Mol Psychiatry. avr 2020;25(4):719-31.

20. Moreno-Ramos OA, Olivares AM, Haider NB, De Autismo LC, Lattig MC. Whole-Exome Sequencing in a South American Cohort Links ALDH1A3, FOXN1 and Retinoic Acid Regulation Pathways to Autism Spectrum Disorders. Schubert M, éditeur. PLoS ONE. 9 sept 2015;10(9):e0135927.

21. Guo M, Zhu J, Yang T, Lai X, Lei Y, Chen J, et al. Vitamin A and vitamin D deficiencies exacerbate symptoms in children with autism spectrum disorders. Nutritional Neuroscience. 2 sept 2019;22(9):637-47.

22. Lerner V, Miodownik C, Gibel A, Kovalyonok E, Shleifer T, Goodman AB, et al. Bexarotene as Add-On to Antipsychotic Treatment in Schizophrenia Patients: A Pilot Open-Label Trial. 2008;31(1).

23. Lerner V, Miodownik C, Gibel A, Sirota P, Bush I, Elliot H, et al. The Retinoid X Receptor Agonist Bexarotene Relieves Positive Symptoms of Schizophrenia: A 6-Week, Randomized, Double-Blind, Placebo-Controlled Multicenter Trial. J Clin Psychiatry. 15 déc 2013;74(12):18230.

24. Castro DS, Hermanson E, Joseph B, Wallén A, Aarnisalo P, Heller A, et al. Induction of cell cycle arrest and morphological differentiation by Nurr1 and retinoids in dopamine MN9D cells. J Biol Chem. 16 nov 2001;276(46):43277-84.

25. Korecka JA, van Kesteren RE, Blaas E, Spitzer SO, Kamstra JH, Smit AB, et al. Phenotypic characterization of retinoic acid differentiated SH-SY5Y cells by transcriptional profiling. PLoS One. 2013;8(5):e63862.

26. Rataj-Baniowska M, Niewiadomska-Cimicka A, Paschaki M, Szyszka-Niagolov M, Carramolino L, Torres M, et al. Retinoic Acid Receptor β Controls Development of Striatonigral Projection Neurons through FGF-Dependent and Meis1-Dependent Mechanisms. J Neurosci. 28 oct 2015;35(43):14467-75.

27. Podleśny-Drabiniok A, Sobska J, De Lera AR, Gołembiowska K, Kamińska K, Dollé P, et al. Distinct retinoic acid receptor (RAR) isotypes control differentiation of embryonal carcinoma cells to dopaminergic or striatopallidal medium spiny neurons. Sci Rep. 20 oct 2017;7(1):13671.

28. Chatzi C, Brade T, Duester G. Retinoic Acid Functions as a Key GABAergic Differentiation Signal in the Basal Ganglia. Palmer TD, éditeur. PLoS Biol. 12 avr 2011;9(4):e1000609.

29. Lane MA, Bailey SJ. Role of retinoid signalling in the adult brain. Progress in Neurobiology. mars 2005;75(4):275-93.

30. Blomhoff R, Blomhoff HK. Overview of retinoid metabolism and function. J Neurobiol. juin 2006;66(7):606-30.

31. Kreżel W, Kastner P, Chambon P. Differential expression of retinoid receptors in the adult mouse central nervous system. Neuroscience. avr 1999;89(4):1291-300.

32. Zetterström RH, Lindqvist E, De Urquiza AM, Tomac A, Eriksson U, Perlmann T, et al. Role of retinoids in the CNS: differential expression of retinoid binding proteins and receptors and evidence for presence of retinoic acid. Eur J of Neuroscience. févr 1999;11(2):407-16.

33. Krȩżel W, Ghyselinck N, Samad TA, Dupé V, Kastner P, Borrelli E, et al. Impaired Locomotion and Dopamine Signaling in Retinoid Receptor Mutant Mice. Science, New Series. 1998;279(5352):863-7.

34. Molotkova N, Molotkov A, Duester G. Role of retinoic acid during forebrain development begins late when Raldh3 generates retinoic acid in the ventral subventricular zone. Developmental Biology. mars 2007;303(2):601-10.

35. Krzyżosiak A, Szyszka-Niagolov M, Wietrzych M, Gobaille S, Muramatsu S ichi, Krężel W. Retinoid X Receptor Gamma Control of Affective Behaviors Involves Dopaminergic Signaling in Mice. Neuron. juin 2010;66(6):908-20.

36. Niewiadomska-Cimicka A, Krzyżosiak A, Ye T, Podleśny-Drabiniok A, Dembélé D, Dollé P, et al. Genome-wide Analysis of RARβ Transcriptional Targets in Mouse Striatum Links Retinoic Acid Signaling with Huntington’s Disease and Other Neurodegenerative Disorders. Mol Neurobiol. juill 2017;54(5):3859-78.

37. Zhang Y, Crofton EJ, Smith TES, Koshy S, Li D, Green TA. Manipulation of retinoic acid signaling in the nucleus accumbens shell alters rat emotional behavior. Behavioural Brain Research. déc 2019;376:112177.

38. Godino A, Salery M, Durand-de Cuttoli R, Estill MS, Holt LM, Futamura R, et al. Transcriptional control of nucleus accumbens neuronal excitability by retinoid X receptor alpha tunes sensitivity to drug rewards. Neuron. mars 2023;S0896627323001174.

39. Marie A, Leroy J, Darricau M, Alfos S, De Smedt-Peyrusse V, Richard E, et al. Preventive Vitamin A Supplementation Improves Striatal Function in 6-Hydroxydopamine Hemiparkinsonian Rats. Front Nutr. 1 févr 2022;9:811843.

40. Paxinos G, Franklin KBJ. The Mouse Brain in Stereotaxic Coordinates.

41. Borel P, Troadec R, Damiani M, Halimi C, Nowicki M, Guichard P, et al. β- Carotene Bioavailability and Conversion Efficiency Are Significantly Affected by Sex in Rats: First Observation Suggesting a Possible Hormetic Regulation of Vitamin A Metabolism in Female Rats. Molecular Nutrition Food Res. nov 2021;65(22):2100650.

42. Marie A, Kinet R, Helbling J, Darricau M, Alfos S, Di Miceli M, et al. Impact of dietary vitamin A on striatal function in adult rats. The FASEB Journal. août 2023;37(8):e23037.

43. Trifilieff P, Martinez D. Imaging addiction: D2 receptors and dopamine signaling in the striatum as biomarkers for impulsivity. Neuropharmacology. janv 2014;76(0 0):498-509.

44. Steel P. The nature of procrastination: a meta-analytic and theoretical review of quintessential self-regulatory failure. Psychol Bull. janv 2007;133(1):65-94.

45. Steel P, König CJ. Integrating Theories of Motivation. AMR. oct 2006;31(4):889-913.

46. Wypych M, Matuszewski J, Dragan WŁ. Roles of Impulsivity, Motivation, and Emotion Regulation in Procrastination – Path Analysis and Comparison Between Students and Non-students. Front Psychol. 5 juin 2018;9:891.

47. Petitet P, Scholl J, Attaallah B, Drew D, Manohar S, Husain M. The relationship between apathy and impulsivity in large population samples. Sci Rep. 1 mars 2021;11(1):4830.

48. Salamone JD, Correa M. The Mysterious Motivational Functions of Mesolimbic Dopamine. Neuron. nov 2012;76(3):470-85.

49. Berke JD. What does dopamine mean? Nat Neurosci. juin 2018;21(6):787-93.

50. Cagniard B, Balsam PD, Brunner D, Zhuang X. Mice with Chronically Elevated Dopamine Exhibit Enhanced Motivation, but not Learning, for a Food Reward. Neuropsychopharmacol. juill 2006;31(7):1362-70.

51. Balci F, Ludvig EA, Abner R, Zhuang X, Poon P, Brunner D. Motivational effects on interval timing in dopamine transporter (DAT) knockdown mice. Brain Research. avr 2010;1325:89-99.

52. Milienne-Petiot M, Kesby JP, Graves M, Van Enkhuizen J, Semenova S, Minassian A, et al. The effects of reduced dopamine transporter function and chronic lithium on motivation, probabilistic learning, and neurochemistry in mice: Modeling bipolar mania. Neuropharmacology. févr 2017;113:260-70.

53. Carpenter AC, Saborido TP, Stanwood GD. Development of hyperactivity and anxiety responses in dopamine transporter-deficient mice. Dev Neurosci. 2012;34(2-3):250-7.

54. Davis GL, Stewart A, Stanwood GD, Gowrishankar R, Hahn MK, Blakely RD. Functional coding variation in the presynaptic dopamine transporter associated with neuropsychiatric disorders drives enhanced motivation and context-dependent impulsivity in mice. Behavioural Brain Research. janv 2018;337:61-9.

55. Yamashita M, Sakakibara Y, Hall FS, Numachi Y, Yoshida S, Kobayashi H, et al. Impaired cliff avoidance reaction in dopamine transporter knockout mice. Psychopharmacology. juin 2013;227(4):741-9.

56. Cinque S, Zoratto F, Poleggi A, Leo D, Cerniglia L, Cimino S, et al. Behavioral Phenotyping of Dopamine Transporter Knockout Rats: Compulsive Traits, Motor Stereotypies, and Anhedonia. Front Psychiatry. 22 févr 2018;9:43.

57. Salahpour A, Medvedev IO, Beaulieu JM, Gainetdinov RR, Caron MG. Local knockdown of genes in the brain using small interfering RNA: a phenotypic comparison with knockout animals. Biol Psychiatry. 1 janv 2007;61(1):65-9.

58. Cagniard B, Sotnikova TD, Gainetdinov RR, Zhuang X. The dopamine transporter expression level differentially affects responses to cocaine and amphetamine. J Neurogenet. 2014;28(1-2):112-21.

59. Zhuang X, Oosting RS, Jones SR, Gainetdinov RR, Miller GW, Caron MG, et al. Hyperactivity and impaired response habituation in hyperdopaminergic mice. Proc Natl Acad Sci U S A. 13 févr 2001;98(4):1982-7.

60. Sotnikova TD, Efimova EV, Gainetdinov RR. Enhanced Dopamine Transmission and Hyperactivity in the Dopamine Transporter Heterozygous Mice Lacking the D3 Dopamine Receptor. IJMS. 3 nov 2020;21(21):8216.

61. Mulvihill KG. Presynaptic regulation of dopamine release: Role of the DAT and VMAT2 transporters. Neurochemistry International. janv 2019;122:94-105.

62. Sánchez-Hernández D, Poon AN, Kubant R, Kim H, Huot PSP, Cho CE, et al. High vitamin A intake during pregnancy modifies dopaminergic reward system and decreases preference for sucrose in Wistar rat offspring. The Journal of Nutritional Biochemistry. janv 2016;27:104-11.

63. Samad TA, Krezel W, Chambon P, Borrelli E. Regulation of dopaminergic pathways by retinoids: Activation of the D2 receptor promoter by members of the retinoic acid receptor–retinoid X receptor family. Proc Natl Acad Sci USA. 23 déc 1997;94(26):14349-54.

64. Liao WL, Tsai HC, Wang HF, Chang J, Lu KM, Wu HL, et al. Modular patterning of structure and function of the striatum by retinoid receptor signaling. Proc Natl Acad Sci USA. 6 mai 2008;105(18):6765-70.

65. Song R, Zhang HY, Li X, Bi GH, Gardner EL, Xi ZX. Increased vulnerability to cocaine in mice lacking dopamine D _3_ receptors. Proc Natl Acad Sci USA. 23 oct 2012;109(43):17675-80.

66. Simpson EH, Winiger V, Biezonski DK, Haq I, Kandel ER, Kellendonk C. Selective overexpression of dopamine D3 receptors in the striatum disrupts motivation but not cognition. Biol Psychiatry. 15 nov 2014;76(10):823-31.

67. Trifilieff P, Feng B, Urizar E, Winiger V, Ward RD, Taylor KM, et al. Increasing dopamine D2 receptor expression in the adult nucleus accumbens enhances motivation. Mol Psychiatry. sept 2013;18(9):1025-33.

68. Gallo EF, Meszaros J, Sherman JD, Chohan MO, Teboul E, Choi CS, et al. Accumbens dopamine D2 receptors increase motivation by decreasing inhibitory transmission to the ventral pallidum. Nat Commun. 14 mars 2018;9(1):1086.

69. Buckholtz JW, Treadway MT, Cowan RL, Woodward ND, Li R, Ansari MS, et al. Dopaminergic network differences in human impulsivity. Science. 30 juill 2010;329(5991):532.

70. Moreno M, Economidou D, Mar AC, López-Granero C, Caprioli D, Theobald DE, et al. Divergent effects of D₂/₃ receptor activation in the nucleus accumbens core and shell on impulsivity and locomotor activity in high and low impulsive rats. Psychopharmacology (Berl). juill 2013;228(1):19-30.

71. Besson M, Belin D, McNamara R, Theobald DE, Castel A, Beckett VL, et al. Dissociable control of impulsivity in rats by dopamine d2/3 receptors in the core and shell subregions of the nucleus accumbens. Neuropsychopharmacology. janv 2010;35(2):560-9.

72. Trifilieff P, Ducrocq F, van der Veldt S, Martinez D. Blunted Dopamine Transmission in Addiction: Potential Mechanisms and Implications for Behavior. Semin Nucl Med. janv 2017;47(1):64-74.

73. Enriquez-Traba J, Arenivar M, Yarur-Castillo HE, Noh C, Flores RJ, Weil T, et al. Dissociable control of motivation and reinforcement by distinct ventral striatal dopamine receptors. Nat Neurosci. janv 2025;28(1):105-21.

74. Veenvliet JV, Smidt MP. Molecular mechanisms of dopaminergic subset specification: fundamental aspects and clinical perspectives. Cell Mol Life Sci. déc 2014;71(24):4703-27.

75. Meng SZ, Ozawa Y, Itoh M, Takashima S. Developmental and age-related changes of dopamine transporter, and dopamine D1 and D2 receptors in human basal ganglia. Brain Res. 2 oct 1999;843(1-2):136-44.

76. Green AL, Eid A, Zhan L, Zarbl H, Guo GL, Richardson JR. Epigenetic Regulation of the Ontogenic Expression of the Dopamine Transporter. Front Genet [Internet]. 4 nov 2019 [cité 8 juill 2025];10. Disponible sur: https://www.frontiersin.org/journals/genetics/articles/10.3389/fgene.2019.01099/fu ll

77. Rao A, Richards TL, Simmons D, Zahniser NR, Sorkin A. Epitope-tagged dopamine transporter knock-in mice reveal rapid endocytic trafficking and filopodia targeting of the transporter in dopaminergic axons. FASEB J. mai 2012;26(5):1921-33.

78. Weinstein JJ, Chohan MO, Slifstein M, Kegeles LS, Moore H, Abi-Dargham A. Pathway-Specific Dopamine Abnormalities in Schizophrenia. Biol Psychiatry. 1 janv 2017;81(1):31-42.

79. Kegeles LS, Abi-Dargham A, Frankle WG, Gil R, Cooper TB, Slifstein M, et al. Increased synaptic dopamine function in associative regions of the striatum in schizophrenia. Arch Gen Psychiatry. mars 2010;67(3):231-9.

80. van Enkhuizen J, Henry BL, Minassian A, Perry W, Milienne-Petiot M, Higa KK, et al. Reduced Dopamine Transporter Functioning Induces High-Reward Risk-Preference Consistent with Bipolar Disorder. Neuropsychopharmacology. déc 2014;39(13):3112-22.

81. Wickens MM, Bangasser DA, Briand LA. Sex Differences in Psychiatric Disease: A Focus on the Glutamate System. Front Mol Neurosci. 5 juin 2018;11:197.

82. Castle DJ, Wessely S, Murray RM. Sex and schizophrenia: effects of diagnostic stringency, and associations with and premorbid variables. Br J Psychiatry. mai 1993;162:658-64.

83. Abel KM, Drake R, Goldstein JM. Sex differences in schizophrenia. Int Rev Psychiatry. 2010;22(5):417-28.

84. Cuffe SP, Moore CG, McKeown RE. Prevalence and correlates of ADHD symptoms in the national health interview survey. J Atten Disord. nov 2005;9(2):392-401.

85. Green T, Flash S, Reiss AL. Sex differences in psychiatric disorders: what we can learn from sex chromosome aneuploidies. Neuropsychopharmacol. janv 2019;44(1):9-21.

86. Markham JA. Sex steroids and schizophrenia. Rev Endocr Metab Disord. sept 2012;13(3):187-207.

87. Hwang WJ, Lee TY, Kim NS, Kwon JS. The Role of Estrogen Receptors and Their Signaling across Psychiatric Disorders. IJMS. 31 déc 2020;22(1):373.

88. Mu E, Gurvich C, Kulkarni J. Estrogen and psychosis — a review and future directions. Arch Womens Ment Health [Internet]. 15 janv 2024 [cité 27 août 2024]; Disponible sur: https://link.springer.com/10.1007/s00737-023-01409-x

