## Supplementary information for "Developmental vitamin A deficiency induces sex-specific reward processing alterations through a dysregulation of the mesolimbic dopamine transmission in mice"

*Couty et al.*

**Contents:**

Supplementary methods

**Supplementary Table 1**. Primer sequences used for gene amplification in RT-qPCR experiments. Genes in italic are housekeeping genes.

**Supplementary Table 2.** Key resources table. (see separate Excel file)

**Supplementary Table 3**. Summary of statistical analysis. (see separate Excel file)

**Supplementary Figure 1**. Developmental VAD alters reward processing in male offspring.

**Supplementary Figure 2**. Effects of developmental VAD on expression of key markers of DA transmission in the striatum and the VTA.

**Supplementary Figure 3**. Developmental VAD potentiates DA dynamics in the NAc of male offspring.

**Supplementary Figure 4**. The pro-motivational effect of amphetamine is blunted in VAD male offspring.

#### SUPPLEMENTARY METHODS

##### Behavioral experiments

**Motivation assessment**.

*Pavlovian training.* During 3 sessions (30 min) the mice were accustomed to the operant box. The levers are retracted, and the reward was delivered in the food port at 2-min intervals on average. The reward was indicated by a brief tone and the illumination of a cue light above the lever. The reward had to be consumed by the animal for a new reward to be delivered.

*Concurrent lever pressing/free-feeding task:* In this effort-based choice task, mice had the choice between lever pressing for the palatable reward under a RR80 schedule or consuming the freely available, less palatable, regular chow (respectively CD or VAD).

**Impulsivity assessment**.

The cages are configured in a similar way, except that this time the two levers are considered "reinforced", associated with the delivery of one reward (low reward lever, LR) for the first lever and three rewards for the second (high reward lever, HR).

*Operant and forced-choice training*: in this first phase, the mice were trained to associate a first lever with the release of the low/single reward. Each lever (right or left) delivering the low reward was randomly assigned to each mouse. Only one lever is presented at the start of each trial, associated with a light stimulus above it which signals the opportunity to obtain a reward. The mice had 30 seconds to press the lever, causing it to retract and extinguish the light and obtain the reward. If the animal does not press the lever within the allotted time, no reward is delivered, and the lever retracts, counting as an omission. After 30 seconds, the lever is available again. The session is over when the animal has pressed the lever 30 times. Contingency between instrumental response and reward attainment is considered acquired when the mice reach a criterion of ≥80% of reinforced presses on all 30 trials for three consecutive days (i.e., three sessions). Once this criterion was reached, the mice were then trained to associate the second lever with the release of the high reward, maintaining the same learning criterion.

*Discrimination and free-choice training*: the second phase give access to the two reinforced levers at the same time. The first 10 trials were "forced", i.e., the two levers were alternately presented to enable the animal to remember the association of the two levers with the number of rewards they dispense. The mice were then free to choose between the two levers during the last 20 trials. The task is considered complete when the animal achieves a preference ≥80% for the lever associated with the high reward on the 20 free trials for three consecutive days.

*Delayed discounting*: this last phase of temporal devaluation involves the progressive increase of a delay of varying length, before the release of the high reward, while the small reward remains available immediately. The delays (2, 4, 6, 8 and 10 seconds), before the release of the high reward following lever pressing, were imposed in separate sessions (3 sessions for each delay). A high preference ratio for each delay was calculated as follows: $\frac{\% \ of \ HR \ lever \ presses \ with \ delay}{\% \ of \ HR \ lever \ presses \ without \ delay}$.

**Outcome devaluation procedure.** Animals were given access to *ad libitum* chow (CD or VAD) or milk reward pellets for 90 minutes prior to the test. They were then placed in the operant chamber under a RR80 schedule of reinforcement. There were at least two outcome devaluation procedures where the prefeeding chow or pellets were counterbalanced between sessions.

**Hedonic reactivity measurement.** The sucrose preference test was used to measure the animal's ability to experience the pleasure of drinking sweetened water, based on the mice's innate preference for sugar. The mice were habituated to the presence of two bottles: one containing water, the other containing a 1% sucrose solution (S7903, Sigma). Bottle position (right or left) was determined randomly to avoid a preference bias for bottle position. The quantities of water or sucrose consumed were measured by weighing the bottles at 24h. A sucrose preference index was calculated as follows: : $\frac{Sucrose \ consumed \ (g)}{Water + \ sucrose \ consumed \ (g)} x100$.

**Locomotion assessment.** An open field was used to analyze spontaneous locomotor and exploration activity. The square-shaped device (40 x 40 cm), surrounded by walls (16 cm high), was divided into two parts: a central zone and a peripheral zone. Each mouse was placed in the central zone and was free to explore the new environment for 10 min. To encourage exploration, the light intensity in the center of the device was relatively low (50 lux). Total distance travelled was recorded using the SMART 2 video tracking system (SMART software, Bioseb). Between each mouse, the arena was cleaned with a 30% ethanol solution.

**Histology**

After washes with PBS 1X (pH=7.4), sections were blocked with a blocking solution (3% donkey serum, 3% bovine serum albumin, 0.3% Triton X-100, PBS 1X), for 1h at room temperature. Sections were then incubated with the primary antibodies diluted in blocking solution overnight at 4°C. The next days, sections were washed multiple times with PBS 1X and incubated with the secondary antibodies conjugated to a fluorochrome for 2h at room temperature, protected from light. Sections were washed multiple times with PBS 1X and then mounted in Vectashield Hardest mounting medium with DAPI (H-1500, VectorLab, Eurobio Scientific). The antibodies used are summarized in Table S2. All sections were scanned using a Nanozoomer slide scanner (Hamamatsu Nanozoomer 2.0 HT) with a 20X objective (20X, NA 0.75). Setting parameters for acquisition were kept constant between all animals for a given labelling. The digital images obtained were processed with QuPath 0.5.0 (University of Edinburgh) with semi-automated quantification to limit analysis bias.

**Tyrosine hydroxylase immunostaining**. For tyrosine hydroxylase (TH) staining in the striatum, intensity of staining was quantified. For each slide, several substructures of the striatum (core, medial shell (mShell) and lateral shell (lShell)) were delineated by hand with the polygon tool, which defines the region of interest (ROI). Staining intensity for TH was measured based on this ROI using the “add intensity features” function, with pixel size at 2 µm and tile diameter at 25 µm. TH-positive (TH+) neurons in the VTA were detected with the function “cell detection” with a request pixel size of 2 µm, a background radius of 30 µm, a sigma of 1.5 µm, and a threshold of 10. The number of TH+ neurons was normalized by the total area ($\mu m^2$), as defined by boundaries.

**Viral infection and fiber implantation site**. For dLight expression in NAc and DREADD Gi expression in VTA dopaminergic neurons, the infection sites were assessed in the targeted structures according to GFP or mCherry fluorescence, respectively.

**Statistical analysis**

Data sets were tested for normality, and subsequent parametric or non-parametric tests were performed accordingly. Unpaired t-tests or Mann-Whitney tests were used for the comparison of two independent groups. For the analysis of more than two groups with one factor, one-way analysis of variance (ANOVA) or the non-parametric Kruskal-Wallis test was used. When two of three factors were present, two-way or three-way ANOVA (repeated measures when necessary) was performed, respectively. The ANOVA was followed by Tukey's post-hoc test and the Kruskal-Wallis test by Dunn's multiple comparison test when applicable. A one-sample t-test was used to compare group means to a fixed value. Correlation between data were analyzed using the Pearson correlation and linear regression.

**Supplementary Table 1. Primer sequences used for gene amplification in RT-qPCR experiments. Genes in italic are housekeeping genes.** RAR: retinoic acid receptor, RXR: retinoid x receptor, DR: dopamine receptor, TH: tyrosine hydroxylase, DDC: dopamine decarboxylase, MAOA: monoamine oxidase A, DAT: dopamine transporter, VMAT: vesicular monoamine transporter 2, GAPDH: glyceraldehyde 3-phosphate dehydrogenase, B2M: beta-2 microglobulin.

| Gene | Sequence forward primer (5'-3') | Sequence reverse primer (5'-3') |
|---|---|---|
| RARβ | ccgcctgcttggatatcttg | gtgtaaggccatcagagaaagtca |
| RXRγ | agtagccacgaagacatgcc | tctccacgttcatgtcaccg |
| D1R | tgccgctgtcatcaggtttc | ttaaggatggatgccgtggag |
| D2R | acccttcgggggaaaacaac | cctccaacttcagctccaacac |
| D3R | tggcaacggtctggtatgtg | gatggcacagaggttcaggatg |
| TH | atgccaaggacaagctcagg | agaaggggctgggaactttg |
| DDC | ttggctgcattggtttctcc | ttggctgcattggtttctcc |
| MAOA | tgaggtatctgccctgtggttc | ccccaaggaggaccattatctg |
| DAT | tcatggttattgccgggatg | ttccaggtgttgttgcagtgg |
| VMAT | ccgtcgtagttcccatcattcc | tccccgtgaacacagtagagttg |
| *GAPDH* | *gaacatcatccctgcatcca* | *ccagtgagcttcccgttca* |
| *B2M* | *agttaagcatgccagtatggcc* | *tctcgatcccagtagacggtct* |

**Supplementary Table 2. Key resources table.** (see separate Excel file)

**Supplementary Table 3. Summary of statistical analysis for Fig.1 to Fig.S4.** (see separate Excel file)

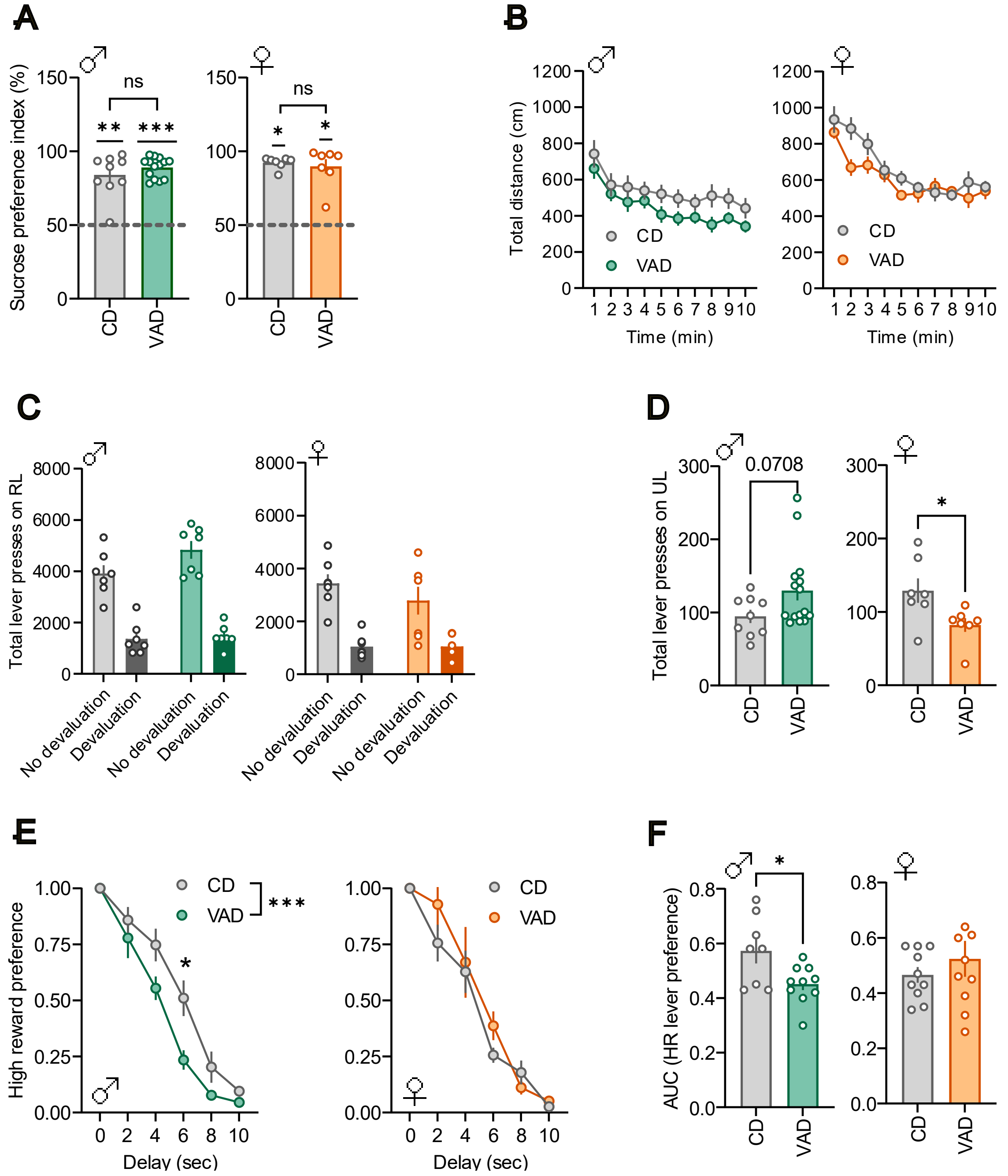

**Supplementary Figure 1 (referent to Figure 1). Developmental VAD alters reward processing in male offspring. (A)** Sucrose preference index (%) in male (Mann-Whitney test: *p=0.5571*; CD: n=9, VAD n=14) and female (Mann-Whitney test: *p=0.5210*; CD: n=7, VAD: n=7) offspring. **(B)** Locomotor activity over time in an open field in male (two-way repeated measure ANOVA: diet *p=0.1697*, time ****$p<0.0001$ and interaction *p=0.7475*; CD: n=9, VAD: n=17) and female offspring (two-way repeated measure ANOVA: diet *p=0.1740*, time ****$p<0.0001$ and interaction *p=0.0716*; CD: n=7, VAD: n=7). **(C)** Total lever presses on the reinforced lever (RL)

during the outcome devaluation procedure in male (two-way ANOVA: diet $p=0.0802$, devaluation ****$p<0.0001$ and interaction $p=0.1485$; CD: n=7, VAD: n=7) and female offspring (two-way ANOVA: diet $p=0.3427$, devaluation ****$p<0.0001$ and interaction $p=0.3322$; CD: n=7, VAD: n=7). **(D)** Total lever presses on the unreinforced lever (UL) during the progressive ratio (PRx2) tasks (Mann-Whitney test: ♂ $p=0.0708$; unpaired t-test: ♀ *$p=0.0312$). **(E)** High reward (HR) preference across the different delays in male (two-way ANOVA: diet ***$p=0.0001$, delay ****$p<0.0001$ and interaction $p=0.1163$; CD: n=8, VAD: n=9) and female (two-way ANOVA: diet $p=0.2332$, delay ****$p<0.0001$ and interaction $p=0.6033$) offspring, in the delay discounting task. **(F)** Area under the curve (AUC) of the HR preference in the delay discounting task in male (unpaired t-test: *$p=0.0209$; CD: n=8, VAD: n=9) and female (Mann-Whitney test: $p=0.5646$; CD: n=10, VAD: n=10) offspring. ANOVA: analysis of variance, CD: control diet, VAD: vitamin A deficient, RL: reinforced lever, UL: unreinforced lever, HR: high reward, AUC: area under the curve. (Table S3 for detailed statistical analysis and results).

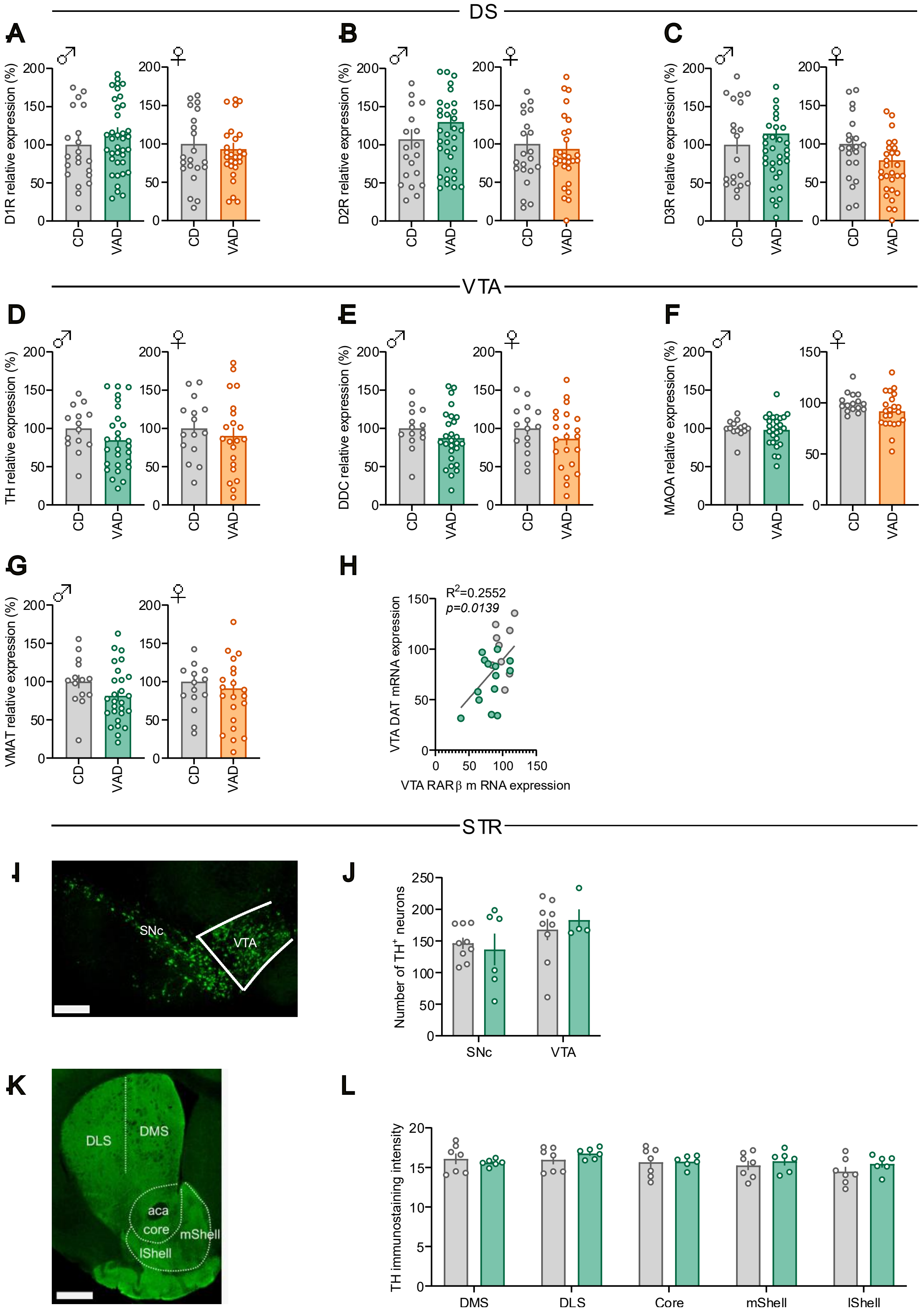
DS
A
D1R relative expression (%)
CD
VAD
B
D2R relative expression (%)
C
D3R relative expression (%)
VTA
D
TH relative expression (%)
E
DDC relative expression (%)
F
MAOA relative expression (%)
G
VMAT relative expression (%)
H
R2=0.2552
p=0.0139
VTA DAT mRNA expression
VTA RARβ m RNA expression
STR
I
SNc
VTA
J
Number of TH+ neurons
K
DLS
DMS
aca
core
mShell
lShell
L
TH immunostaining intensity
DMS
DLS
Core
mShell
lShell

**Supplementary Figure 2 (referent to Figure 2). Effects of developmental VAD on expression of key markers of DA transmission in the striatum and the VTA. (A)** mRNA relative expression of genes coding for D1R (Mann-Whitney test: ♂ *p=0.2020*; ♀ *p=0.9884*), **(B)** D2R (Mann-Whitney test: ♂ *p=0.1673*; ♀ *p=0.9587*) and **(C)** D3R (Mann-Whitney test: ♂ *p=0.6322*; ♀ *p=0.0601*) in the dorsal striatum (DS) in male (CD: n=23, VAD: n=39) and female (CD: n=20, VAD: n=30) offspring. **(D)** mRNA relative expression of genes coding for TH (unpaired t-test: ♂ *p=0.2163*; ♀ *p=0.5332*), **(E)** DDC (unpaired t-test: ♂ *p=0.2495*; ♀ *p=0.2719*), **(F)** MAOA (Mann-Whitney test: ♂ *p=0.8599*; unpaired t-test: ♀ *p=0.1011*) and **(G)** VMAT (unpaired t-test: ♂ *p=0.1206*; Mann-Whitney test: ♀ *p=0.6211*) in the ventral tegmental area (VTA) in male (CD: n=24, VAD: n=27) and female (CD: n=15, VAD: n=22) offspring. (**H**) Individual correlation between mRNA expression of RARβ and DAT in the VTA in males ($R^2$=0.2552; *p=0.0139;* CD: n=9, VAD n=14). **(I)** Representative image of the midbrain immunostained for TH (scale bar: 250µm). **(J)** Quantification of the number of TH+ neurons in males (two-way ANOVA: diet *p=0.2792*, structure *p=0.1485* and interaction *p=0.6723*; CD: n=7, VAD: n=6). **(K)** Representative image of the striatum immunostained for TH (scale bar: 800µm). **(L)** Quantification of fluorescence intensity of TH staining in DS and nucleus accumbens (NAc) in males (two-way ANOVA: diet *p=0.885*, structure *p=0.0703* and interaction *p=0.4976*; CD: n=9, VAD: n=4-6). CD: control diet, VAD: vitamin A deficient, DR: dopamine receptor, TH: tyrosine hydroxylase, DDC: dopamine decarboxylase, MAOA: monoamine oxidase, VMAT: vesicular monoamine transporter, DS: dorsal striatum, NAc: nucleus accumbens, VTA: ventral tegmental area, SNc: substantia nigra *pars compacta,* DMS: dorsomedial striatum, DLS: dorsolateral striatum, mShell: medial shell, lShell: lateral shell, RAR: retinoic acid receptor. (Table S3 for detailed statistical analysis and results).

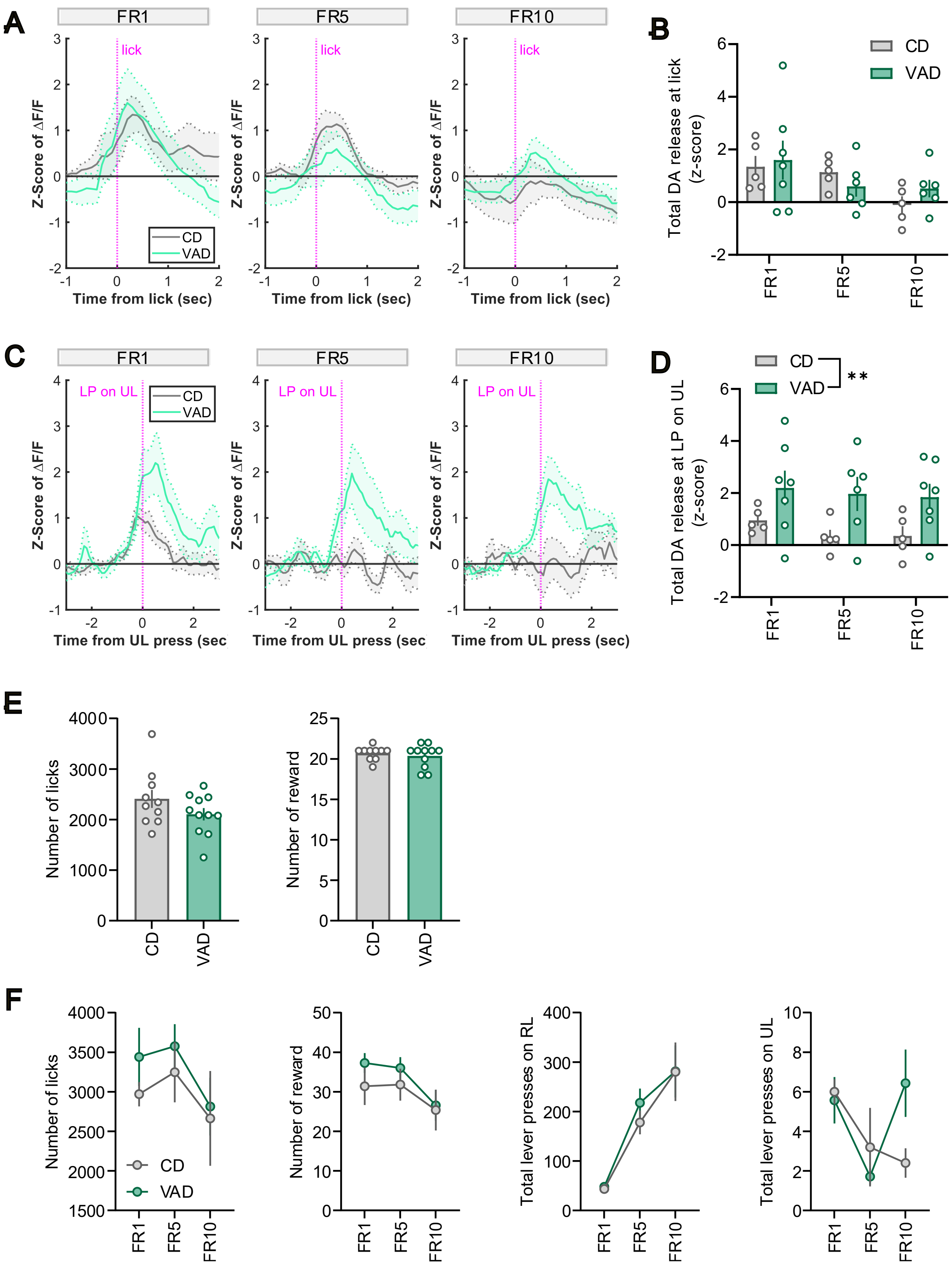

**Supplementary Figure 3 (referent to Figure 3). Developmental VAD potentiates DA dynamics in the NAc of male offspring. (A)** Dopamine (DA) dynamic (z-score of ΔF/F) during fixed ratio (FR1, FR5, FR10) trials aligned onto the lick onset in males

(CD: n=5, VAD: n=7) and **(B)** quantification of the peak amplitude (z-score) (0 to 2 sec) (two-way ANOVA: diet *p=0.7821*, ratio *p=0.0488* and interaction *p=0.7821*). **(C)** DA dynamic during fixed ratio (FR1, FR5 and FR10) trials aligned onto of the unreinforced lever (UL) press in males (CD: n=5; VAD: n=7) and **(D)** quantification (0 to 2 sec) (two-way ANOVA: diet ***p=0.0024*, ratio *p=0.6182* and interaction *p=0.9300*). **(E)** Quantification of the DA dynamic before the onset of reinforced lever (RL) press (-1 to 0 sec) for the different fixed ratio trials (two-way ANOVA: diet *p=0.1145*, ratio ****$p<0.0001$ and interaction *p=0.6954*) in males (CD: n=5, VAD: n=7). **(F)** Number of licks (Mann-Whitney test: *p=0.4078*) and reward (Mann-Whitney test: *p=0.6970*) during Pavlovian conditioning in male offspring (CD: n=5, VAD: n=7). **(G)** Number of licks (two-way ANOVA: diet *p=0.3162*, ratio *p=0.2085* and interaction *p=0.9142*), reward (two-way ANOVA: diet *p=0.2357*, ratio *p=0.0614* and interaction *p=0.8216*), RL (two-way ANOVA: diet *p=0.6216*, ratio ****$p<0.0001$ and interaction *p=0.8432*) and UL presses (two-way ANOVA: diet *p=0.5039*, ratio **p=0.0452* and interaction *p=0.0886*) during fixed ratio trials (FR1, FR5, FR10) in male offspring (CD: n=5, VAD: n=7). ANOVA: analysis of variance, CD: control diet, VAD: vitamin A deficient, FR: fixed ratio, LP: lever press, RL: reinforced lever, UL: unreinforced lever, DA: dopamine. (Table S3 for detailed statistical analysis and results).

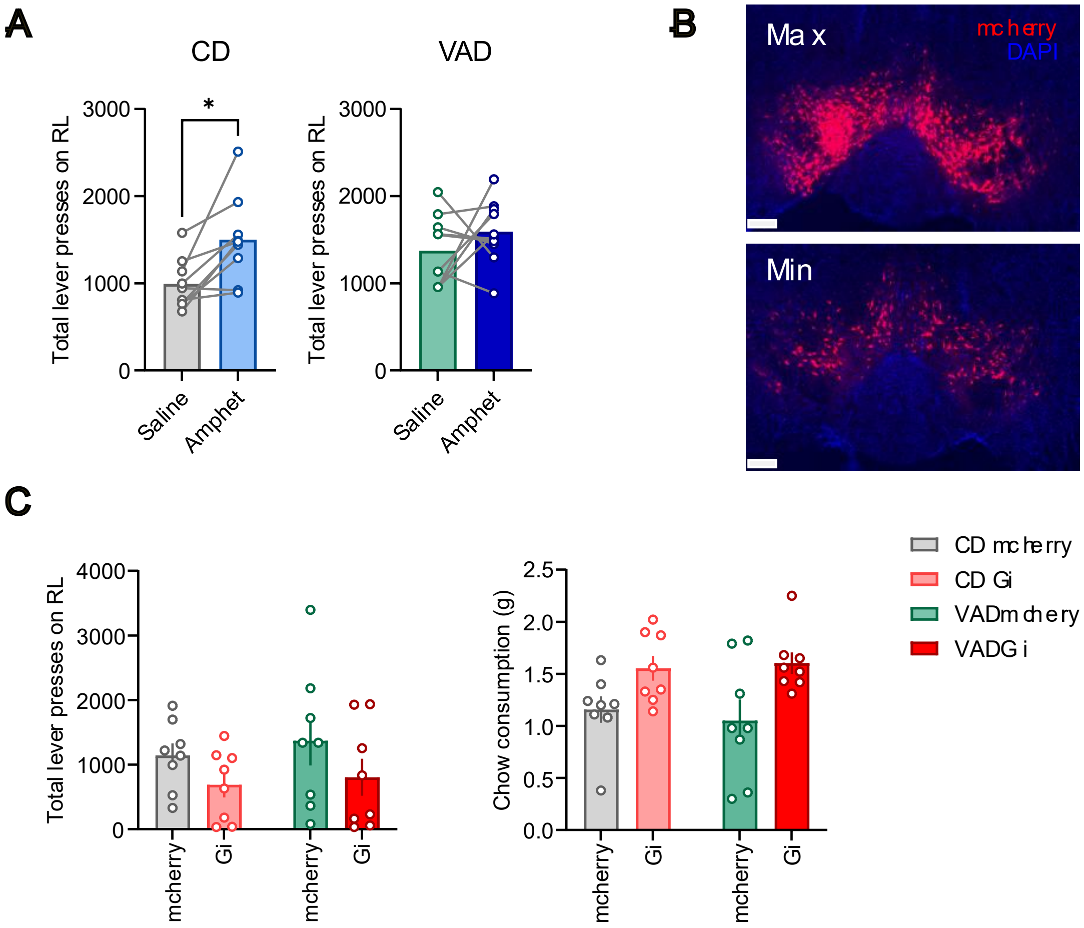

**Supplementary Figure 4 (referent to Figure 4). The pro-motivational effect of amphetamine is blunted in VAD male offspring. (A)**Total lever presses on the reinforced lever (RL) in CD (unpaired t-test: *$p$=*0.0171;* n=9) and VAD (unpaired t-test: p=*0.2136;* n=10) male offspring after saline or amphetamine (amphet) injection. **(B)** Maximal and minimal viral expression in the VTA (scale bar=200µm). **(C)** Total lever presses on the RL (two-way ANOVA: diet *p=0.5340*, virus *p=0.0748* and interaction *p=0.8423*) and **(F)** chow consumption (two-way ANOVA: diet *p=0*.8488, virus ***p=0.0025* and interaction *p=0.5918*) in the concurrent lever pressing/free-feeding task under CNO, in male offspring (mcherry CD: n=8, mcherry VAD: n=8, Gi CD: n=8, Gi VAD: n=8). CD: control diet, VAD: vitamin A deficient diet, RL: reinforced lever. (Table S3 for detailed statistical analysis and results).

### KEY RESOURCES TABLE

| Resource Type | Specific Reagent or Resource | Source or Reference | Identifiers |
|---|---|---|---|
| Antibody | Rabbit anti-DAT (used at 1/1000) | Millipore | AB2231 |
| Antibody | Mouse anti-α-tubulin (used at 1/2000) | Merck | T5168 |
| Antibody | Anti-rabbit IRDye® 800CW (used at 1/10 000) | LICOR | 926-32211 |
| Antibody | Anti-mouse IREDye® 680RD (used at 1/10 000) | LICOR | 926-68071 |
| Antibody | Sheep anti-TH (used at 1/1000) | Millipore | AB1542 |
| Antibody | Anti-sheep IgG Alexa 488 (used at 1/1000) | Abcam | ab175713 |
| Antibody | Chicken anti-mcherry (used at 1/1000) | Abcam | ab205402 |
| Antibody | Anti-chicken IgY Alexa 647 (used at 1/1000) | Abcam | ab150171 |
| Bacterial or Viral Strain | AAV-9-hSyn1-chl-dLight1.3b-WPRE-bGHp(A) | Zurich Vector Facility | v565-9 |
| Bacterial or Viral Strain | AAV8-hSyn1-dlox-HM4D(Gi)-mcherry(rev)-dlox-WPRE-hGHp(A) | Zurich Vector Facility | v84-8 |
| Bacterial or Viral Strain | AAV8-hSyn1-dlox-mcherry(rev)-dlox-WPRE-hGHp(A) | Zurich Vector Facility | v166-8 |
| Biological Sample | N/A | N/A | N/A |
| Cell Line | N/A | N/A | N/A |
| Chemical Compound or Drug | d-(+)-Amphetamine sulfate | LGC Standards | 10-D-18 |
| Chemical Compound or Drug | Clozapine-N-Oxide (CNO) | ENZO | BML-NS105-0025 |
| Chemical Compound or Drug | Sucrose | Sigma | S7903 |
| Commercial Assay or Kit | BC Assay protein quantification kit | Interchim | FT-40840A |
| Deposite Data or Public Database | N/A | N/A | N/A |
| Genetic Reagent | N/A | N/A | N/A |
| Organism / Strain | Mouse : C57BL6/JRj (Male and Female) | Janvier Labs | N/A |
| Organism / Strain | Mouse : DAT$^{IREScre}$, B6.SJL-Slc6a3tm1.1(cre)Bkmn/J | The Jackson Laboratory | RRID: IMSR_JAX:006660 |
| Peptide or Recombinant Protein | N/A | N/A | N/A |
| Recombinant DNA | N/A | N/A | N/A |
| Sequence-Based Reagent | Primers for RT-qPCR, see Table S1 | Eurogentec | N/A |
| Transfected construct | N/A | N/A | N/A |
| Software, Algorithm | GraphPad-Prism 10 | GraphPad Software | RRID: SCR_002798 |
| Software, Algorithm | Multichannel Fiber Photometry Software | RWD | v2.0.0.33169 |
| Software, Algorithm | MATLAB | MathWorks | R2023b |
| Software, Algorithm | LightCycler® 480 Software release 1.5.0 (1.5.0.39) | Roche | N/A |
| Software, Algorithm | GenEx v.7 | Applied Biosystems™ | N/A |
| Software, Algorithm | Image Studio® Software | LICORbio | v. 5.2.5 |
| Software, Algorithm | QuPath | University of Edinburg | v. 0.5.0 |
| Other | Control Diet (7.5 IU of retinol/g of diet) | Safe diets | U8958A01R 00426 |
| Other | Vit. A deficient diet (0.4 IU of retinol/g of diet) | Safe diets | U8958A01R 00425 |
| Other | Rodent Pellet Tablet (20 mg) with non-added vitamin A, non-flavoured | Test Diet | RBUP |

| Figure panels | Outcome measure | n | Statistical analysis | | F-value | p-value | Multiple comparaisons |
|---|---|---|---|---|---|---|---|
| Fig. 1C ♂ | Number of RL presses/min | CD: 9 VAD: 15 | 2-way ANOVA | Diet | F(1,85)=0,1093 | p=0,7418 | |
| | | | | Ratio | F(3,85)=221,1 | **p<0,0001** | |
| | | | | Interaction | F(3,85)=1,434 | p=0,2385 | |
| Fig. 1C ♀ | Number of RL presses/min | CD: 7 VAD: 7 | 2-way ANOVA | Diet | F(1,48)=1,522 | p=0,2234 | |
| | | | | Ratio | F(3,48)=56,67 | **p<0,0001** | |
| | | | | Interaction | F(3,48)=0,2961 | p=0,8280 | |
| Fig. 1E ♂ | Total lever presses on RL (PRx2) | CD: 9 VAD: 15 | Normality test | Anderson-Darling | | p=0,4746 | |
| | | | | D'Agostino-Pearson test | | p=0,0562 | |
| | | | | Shapiro-Wilk | | p=0,3805 | |
| | | | | Kolmogorov-Smirnov | | p>0,0001 | |
| | | | Unpaired t-test | | | **p=0,0039** | |
| Fig. 1E ♀ | Total lever presses on RL(PRx2) | CD: 7 VAD: 7 | Normality test | Anderson-Darling | | p=0,2770 | |
| | | | | D'Agostino-Pearson test | | p=0,4034 | |
| | | | | Shapiro-Wilk | | p=0,1987 | |
| | | | | Kolmogorov-Smirnov | | p>0,0001 | |
| | | | Unpaired t-test | | | p=0,7958 | |
| Fig. 1F ♂ | Breakpoint (PRx2) | CD: 9 VAD: 15 | Normality test | Anderson-Darling | | p=0,0570 | |
| | | | | D'Agostino-Pearson test | | **p=0,0286** | |
| | | | | Shapiro-Wilk | | p=0,0580 | |
| | | | | Kolmogorov-Smirnov | | **p=0,0192** | |
| | | | Mann-Whitney test | | | **p=0,0018** | |
| Fig. 1F ♀ | Breakpoint (PRx2) | CD: 7 VAD: 7 | Normality test | Anderson-Darling | | p=0,4702 | |
| | | | | D'Agostino-Pearson test | | p=0,3442 | |
| | | | | Shapiro-Wilk | | p=0,4940 | |
| | | | | Kolmogorov-Smirnov | | p>0,0001 | |
| | | | Unpaired t-test | | | p=0,7087 | |
| Fig. 1H ♂ | Total lever presses on RL (PRx2) | CD: 9 VAD: 7 P0: 10 P21:9 | Normality test | Anderson-Darling | | p=0,1919 | Tukey's pos-hoc test: CD vs VAD **p=0,0523** CD vs P0 p=0,9903 CD vs P21 p=0,2050 VAD vs P0 **p=0,0238** VAD vs P21 p=0,8819 P0 vs P21 p=0,1079 |
| | | | | D'Agostino-Pearson test | | p=0,5019 | |
| | | | | Shapiro-Wilk | | p=0,2302 | |
| | | | | Kolmogorov-Smirnov | | p=0,1000 | |
| | | | One-way ANOVA | Diet | F(3,30)=4,437 | **p=0,0107** | |
| Fig. 1I ♂ | Breakpoint (PRx2) | CD: 9 VAD: 7 P0: 10 P21:9 | Normality test | Anderson-Darling | | p=0,5167 | Tukey's pos-hoc test: CD vs VAD **p=0,0,399** CD vs P0 p=0,9473 CD vs P21 p=0,5395 VAD vs P0 **p=0,0101** VAD vs P21 p=0,4629 P0 vs P21 p=0,2453 |
| | | | | D'Agostino-Pearson test | | p=0,3972 | |
| | | | | Shapiro-Wilk | | p=0,4457 | |
| | | | | Kolmogorov-Smirnov | | p=0,1000 | |
| | | | One-way ANOVA | Diet | F(3,30)=4,487 | **p=0,0102** | |

| Figure panels | Outcome measure | n | Statistical analysis | | F-value | p-value | Multiple comparaisons |
|---|---|---|---|---|---|---|---|
| Fig. 2A ♂ | Plasma retinol (µmolL) | CD: 8<br>VAD: 12 | Normality test | Anderson-Darling | | p=0,4814 | |
| | | | | D'Agostino-Pearson test | | p=0,5536 | |
| | | | | Shapiro-Wilk | | p=0,6881 | |
| | | | | Kolmogorov-Smirnov | | p>0,0001 | |
| | | | Unpaired t-test | | | **p<0,0001** | |
| Fig. 2A ♀ | Plasma retinol (µmolL) | CD: 6<br>VAD: 8 | Normality test | Anderson-Darling | | p=0,6048 | |
| | | | | D'Agostino-Pearson test | | p=0,9295 | |
| | | | | Shapiro-Wilk | | p=0,7637 | |
| | | | | Kolmogorov-Smirnov | | p>0,0001 | |
| | | | Unpaired t-test | | | **p=0,0069** | |
| Fig. 2B ♂ | Hepatic retinol (nmol/g of liver) | CD: 14<br>VAD: 25 | Normality test | Anderson-Darling | | **p<0,0001** | |
| | | | | D'Agostino-Pearson test | | **p<0,0001** | |
| | | | | Shapiro-Wilk | | **p<0,0001** | |
| | | | | Kolmogorov-Smirnov | | **p<0,0001** | |
| | | | Mann-Whitney test | | | **p<0,0001** | |
| Fig. 2B ♀ | Hepatic retinol (nmol/g of liver) | CD: 11<br>VAD: 13 | Normality test | Anderson-Darling | | **p=0,0023** | |
| | | | | D'Agostino-Pearson test | | p=0,0715 | |
| | | | | Shapiro-Wilk | | **p=0,0169** | |
| | | | | Kolmogorov-Smirnov | | **p=0,0017** | |
| | | | Mann-Whitney test | | | **p<0,0001** | |
| Fig. 2C ♂ | Hepatic retinyl esters (nmol/g of liver) | CD: 14<br>VAD: 25 | Normality test | Anderson-Darling | | **p<0,0001** | |
| | | | | D'Agostino-Pearson test | | **p<0,0001** | |
| | | | | Shapiro-Wilk | | **p<0,0001** | |
| | | | | Kolmogorov-Smirnov | | **p<0,0001** | |
| | | | Mann-Whitney test | | | **p<0,0001** | |
| Fig. 2C ♀ | Hepatic retinyl esters (nmol/g of liver) | CD: 11<br>VAD: 13 | Normality test | Anderson-Darling | | **p=0,0141** | |
| | | | | D'Agostino-Pearson test | | p=0,4980 | |
| | | | | Shapiro-Wilk | | p=0,0930 | |
| | | | | Kolmogorov-Smirnov | | **p=0,0069** | |
| | | | Mann-Whitney test | | | **p<0,0001** | |
| Fig. 2D ♂ | DS RARβ relative expression (%) | CD: 14<br>VAD: 25 | Normality test | Anderson-Darling | | p=0,7800 | |
| | | | | D'Agostino-Pearson test | | p=0,8802 | |
| | | | | Shapiro-Wilk | | p=0,9524 | |
| | | | | Kolmogorov-Smirnov | | p>0,0001 | |
| | | | Unpaired t-test | | | p=0,7017 | |
| Fig. 2D ♀ | DS RARβ relative expression (%) | CD: 15<br>VAD: 23 | Normality test | Anderson-Darling | | **p=0,0175** | |
| | | | | D'Agostino-Pearson test | | **p=0,0355** | |
| | | | | Shapiro-Wilk | | **p=0,0396** | |
| | | | | Kolmogorov-Smirnov | | **p=0,0189** | |
| | | | Mann-Whitney test | | | p=0,2043 | |

| | | | | | | |
|---|---|---|---|---|---|---|
| Fig. 2E ♂ | NAc RARβ relative expression (%) | CD: 14<br>VAD: 25 | Normality test | Anderson-Darling | | p=0,5607 |
| | | | | D'Agostino-Pearson test | | p=0,4248 |
| | | | | Shapiro-Wilk | | p=0,5356 |
| | | | | Kolmogorov-Smirnov | | p>0,0001 |
| | | | Unpaired t-test | | | **p=0,0058** |
| Fig. 2E ♀ | NAc RARβ relative expression (%) | CD: 15<br>VAD: 23 | Normality test | Anderson-Darling | | p=0,3400 |
| | | | | D'Agostino-Pearson test | | p=0,1403 |
| | | | | Shapiro-Wilk | | p=0,3652 |
| | | | | Kolmogorov-Smirnov | | p>0,0001 |
| | | | Unpaired t-test | | | **p=0,0006** |
| Fig. 2F ♂ | VTA RARβ relative expression (%) | CD: 14<br>VAD: 27 | Normality test | Anderson-Darling | | p=0,3097 |
| | | | | D'Agostino-Pearson test | | p=0,7368 |
| | | | | Shapiro-Wilk | | p=0,4728 |
| | | | | Kolmogorov-Smirnov | | p>0,0001 |
| | | | Unpaired t-test | | | **p=0,0426** |
| Fig. 2F ♀ | VTA RARβ relative expression (%) | CD: 16<br>VAD: 22 | Normality test | Anderson-Darling | | p=0,0588 |
| | | | | D'Agostino-Pearson test | | p=0,2743 |
| | | | | Shapiro-Wilk | | **p=0,0455** |
| | | | | Kolmogorov-Smirnov | | p=0,0667 |
| | | | Mann-Whitney test | | | **p=0,0009** |
| Fig. 2G ♂ | DS RXRγ relative expression (%) | CD: 14<br>VAD: 25 | Normality test | Anderson-Darling | | p=0,1178 |
| | | | | D'Agostino-Pearson test | | p=0,2080 |
| | | | | Shapiro-Wilk | | p=0,0825 |
| | | | | Kolmogorov-Smirnov | | p>0,0001 |
| | | | Unpaired t-test | | | p=0,1329 |
| Fig. 2G ♀ | DS RXRγ relative expression (%) | CD: 15<br>VAD: 23 | Normality test | Anderson-Darling | | **p=0,0086** |
| | | | | D'Agostino-Pearson test | | **p=0,0039** |
| | | | | Shapiro-Wilk | | **p=0,0075** |
| | | | | Kolmogorov-Smirnov | | **p=0,0271** |
| | | | Mann-Whitney test | | | p=0,0802 |
| Fig. 2H ♂ | NAc RXRγ relative expression (%) | CD: 14<br>VAD: 25 | Normality test | Anderson-Darling | | p=0,7530 |
| | | | | D'Agostino-Pearson test | | p=0,6927 |
| | | | | Shapiro-Wilk | | p=0,8948 |
| | | | | Kolmogorov-Smirnov | | p>0,0001 |
| | | | Unpaired t-test | | | **p=0,0012** |
| Fig. 2H ♀ | NAc RXRγ relative expression (%) | CD: 15<br>VAD: 23 | Normality test | Anderson-Darling | | p=0,3157 |
| | | | | D'Agostino-Pearson test | | p=0,1916 |
| | | | | Shapiro-Wilk | | p=0,4274 |
| | | | | Kolmogorov-Smirnov | | p>0,0001 |
| | | | Unpaired t-test | | | **p<0,0001** |

| | | | | | | |
|---|---|---|---|---|---|---|
| Fig. 2I ♂ | VTA RXRγ relative expression (%) | CD: 14<br>VAD: 27 | Normality test | Anderson-Darling | | p=0,7717 |
| | | | | D'Agostino-Pearson test | | p=0,7938 |
| | | | | Shapiro-Wilk | | p=0,9215 |
| | | | | Kolmogorov-Smirnov | | p>0,0001 |
| | | | Unpaired t-test | | | p=0,7396 |
| Fig. 2I ♀ | VTA RXRγ relative expression (%) | CD: 16<br>VAD: 22 | Normality test | Anderson-Darling | | **p=0,0105** |
| | | | | D'Agostino-Pearson test | | p=0,1558 |
| | | | | Shapiro-Wilk | | **p=0,0133** |
| | | | | Kolmogorov-Smirnov | | **p=0,0049** |
| | | | Unpaired t-test | | | p=0,1808 |
| Fig. 2J ♂ | NAc D1R relative expression (%) | CD: 23<br>VAD: 39 | Normality test | Anderson-Darling | | p=0,3392 |
| | | | | D'Agostino-Pearson test | | p=0,7558 |
| | | | | Shapiro-Wilk | | p=0,5251 |
| | | | | Kolmogorov-Smirnov | | p>0,0001 |
| | | | Unpaired t-test | | | p=0,1501 |
| Fig. 2J ♀ | NAc D1R relative expression (%) | CD: 20<br>VAD: 30 | Normality test | Anderson-Darling | | p=0,3817 |
| | | | | D'Agostino-Pearson test | | p=0,5183 |
| | | | | Shapiro-Wilk | | p=0,4684 |
| | | | | Kolmogorov-Smirnov | | p>0,0001 |
| | | | Unpaired t-test | | | **p=0,0045** |
| Fig. 2K ♂ | NAc D2R relative expression (%) | CD: 23<br>VAD: 39 | Normality test | Anderson-Darling | | p=0,3806 |
| | | | | D'Agostino-Pearson test | | p=0,5194 |
| | | | | Shapiro-Wilk | | p=0,3041 |
| | | | | Kolmogorov-Smirnov | | p>0,0001 |
| | | | Unpaired t-test | | | p=0,8520 |
| Fig. 2K ♀ | NAc D2R relative expression (%) | CD: 20<br>VAD: 30 | Normality test | Anderson-Darling | | p=0,8937 |
| | | | | D'Agostino-Pearson test | | p=0,4816 |
| | | | | Shapiro-Wilk | | p=0,8055 |
| | | | | Kolmogorov-Smirnov | | p>0,0001 |
| | | | Unpaired t-test | | | p=0,0533 |
| Fig. 2L ♂ | NAc D3R relative expression (%) | CD: 23<br>VAD: 39 | Normality test | Anderson-Darling | | p=0,0559 |
| | | | | D'Agostino-Pearson test | | p=0,0804 |
| | | | | Shapiro-Wilk | | **p=0,0378** |
| | | | | Kolmogorov-Smirnov | | p>0,0001 |
| | | | Man-Whitney test | | | **p=0,0020** |
| Fig. 2L ♀ | NAc D3R relative expression (%) | CD: 20<br>VAD: 30 | Normality test | Anderson-Darling | | **p=0,0067** |
| | | | | D'Agostino-Pearson test | | **p=0,0200** |
| | | | | Shapiro-Wilk | | **p=0,0032** |
| | | | | Kolmogorov-Smirnov | | **p=0,0092** |
| | | | Man-Whitney test | | | **p=0,0003** |

| | | | | | | |
|---|---|---|---|---|---|---|
| Fig. 2M ♂ | VTA DAT relative expression (%) | CD: 14<br>VAD: 27 | Normality test | Anderson-Darling | | p=0,3445 |
| | | | | D'Agostino-Pearson test | | p=0,7035 |
| | | | | Shapiro-Wilk | | p=0,3009 |
| | | | | Kolmogorov-Smirnov | | p>0,0001 |
| | | | Unpaired t-test | | | **p<0,0001** |
| Fig. 2M ♀ | VTA DAT relative expression (%) | CD: 15<br>VAD: 22 | Normality test | Anderson-Darling | | p=0,3870 |
| | | | | D'Agostino-Pearson test | | p=0,4393 |
| | | | | Shapiro-Wilk | | p=0,1251 |
| | | | | Kolmogorov-Smirnov | | p>0,0001 |
| | | | Unpaired t-test | | | p=0,4782 |
| Fig. 2N ♂ | VTA DAT protein relative quantity (%) | CD: 18<br>VAD: 17 | Normality test | Anderson-Darling | | p=0,5706 |
| | | | | D'Agostino-Pearson test | | p=0,6366 |
| | | | | Shapiro-Wilk | | p=0,5438 |
| | | | | Kolmogorov-Smirnov | | p>0,0001 |
| | | | Unpaired t-test | | | **p=0,0154** |
| Fig. 2N ♀ | VTA DAT protein relative quantity (%) | CD: 11<br>VAD: 12 | Normality test | Anderson-Darling | | p=0,7860 |
| | | | | D'Agostino-Pearson test | | p=0,8381 |
| | | | | Shapiro-Wilk | | p=0,6147 |
| | | | | Kolmogorov-Smirnov | | p>0,0001 |
| | | | Unpaired t-test | | | p=0,1865 |

| Figure panels | Outcome measure | n | Statistical analysis | | F-value | p-value | Multiple comparaisons |
|---|---|---|---|---|---|---|---|
| Fig. 3D | CS z-score in Pavlovian | CD: 10<br>VAD: 11 | Normality test | Anderson-Darling | | p=0,2303 | |
| | | | | D'Agostino-Pearson test | | p=0,7324 | |
| | | | | Shapiro-Wilk | | p=0,1901 | |
| | | | | Kolmogorov-Smirnov | | p>0,1000 | |
| | | | Unpaired t-test | | | p=0,9317 | |
| Fig. 3E | Lick z-score in Pavlovian | CD: 10<br>VAD: 11 | Normality test | Anderson-Darling | | **p=0,0117** | |
| | | | | D'Agostino-Pearson test | | p=0,2151 | |
| | | | | Shapiro-Wilk | | **p=0,0146** | |
| | | | | Kolmogorov-Smirnov | | **p=0,0245** | |
| | | | Mann-Whitney test | | | p=0,7564 | |
| Fig. 3G | CS z-score in FR1/FR5/FR10 | CD: 5<br>VAD: 7 | 2-way ANOVA | Diet | F(1,29)=7,839 | **p=0,0090** | |
| | | | | Ratio | F(2,29)=1,053 | p=0,3617 | |
| | | | | Interaction | F(2,29)=0,01548 | p=0,9846 | |
| Fig. 3I | First lever press on RL z-score in FR5/FR10 | CD: 5<br>VAD: 7 | 2-way ANOVA | Diet | F(1,19)=4,386 | **p=0,0499** | |
| | | | | Ratio | F(1,19)=0,9463 | p=0,3429 | |
| | | | | Interaction | F(1,19)=0,03595 | p=0,8516 | |

| Figure panels | Outcome measure | n | Statistical analysis | | F-value | p-value | Multiple comparisons |
|---|---|---|---|---|---|---|---|
| Fig. 4A | Total lever presses on RL ratio amphet/saline | CD: 9<br>VAD: 10 | Normality test | Anderson-Darling | | p=0,658 | |
| | | | | D'Agostino-Pearson test | | p=0,3271 | |
| | | | | Shapiro-Wilk | | p=0,0887 | |
| | | | | Kolmogorov-Smirnov | | **p=0,0436** | |
| | | | One sample Wilcoxon test (compared to 1) | | | CD: **p=0,0078**<br>VAD: p=0,3887 | |
| Fig. S5A | Number of RL presses/min (without CNO) | mcherry CD : 8<br>mcherry VAD: 8 Gi CD: 8 Gi VAD: 8 | 3-way ANOVA | Diet | F(1,112)=0,3023 | p=0,5835 | |
| | | | | Virus | F(1,112)=0,008376 | p=0,9272 | |
| | | | | Ratio | F(3,112)=292,9 | **p<0,0001** | |
| | | | | Diet x Virus | F(1,112)=0,4928 | p=0,4842 | |
| | | | | Diet x Ratio | F(3,112)=0,2015 | p=0,8952 | |
| | | | | Ratio x Virus | F(3,112)=0,1539 | p=0,9270 | |
| | | | | Diet x Virus x Ratio | F(3,112)=0,1127 | p=0,9525 | |
| Fig. 4E | Total lever presses on RL (PRx2) under CNO | mcherry CD : 8<br>mcherry VAD: 8 Gi CD: 8 Gi VAD: 8 | 2-way ANOVA | Diet | F(1,28)=0,4579 | p=0,5041 | |
| | | | | Virus | F(1,28)=2,824 | p=0,1040 | |
| | | | | Interaction | F(1,28)=5,454 | **p=0,0269** | mcherry CD vs mcherry VAD: p=0,1684<br>mcherry VAD vs Gi VAD: **p=0,0392** |
| Fig. 4F | Breakpoint (PRx2) under CNO | mcherry CD : 8<br>mcherry VAD: 8 Gi CD: 8 Gi VAD: 8 | 2-way ANOVA | Diet | F(1,28)=3,395 | p=0,0760 | |
| | | | | Virus | F(1,28)=3,395 | p=0,0760 | |
| | | | | Interaction | F(1,28)=5,072 | **p=0,0323** | mcherry CS vs mcherry VAD: **p=0,0345**<br>mcherry VAD vs Gi VAD: **p=0,0345** |

| Figure panels | Outcome measure | n | Statistical analysis | | F-value | p-value | Multiple comparaisons |
|---|---|---|---|---|---|---|---|
| Fig. S1A ♂ | Sucrose preference index (%) | CD: 9<br>VAD: 14 | Normality test | Anderson-Darling | | p=0,0541 | |
| | | | | D'Agostino-Pearson test | | **p=0,0014** | |
| | | | | Shapiro-Wilk | | **p=0,0114** | |
| | | | | Kolmogorov-Smirnov | | p=0,0975 | |
| | | | Mann-Whitney test | | | **p=0,5571** | |
| | | | One sample Wilcoxon test (compared to 50) | | | CD: **p=0,0039**<br>VAD: **p=0,0001** | |
| Fig. S1A ♀ | Sucrose preference index (%) | CD: 7<br>VAD: 7 | Normality test | Anderson-Darling | | **p=0,0072** | |
| | | | | D'Agostino-Pearson test | | **p<0,0001** | |
| | | | | Shapiro-Wilk | | **p=0,0028** | |
| | | | | Kolmogorov-Smirnov | | **p=0,0414** | |
| | | | Mann-Whitney test | | | p=0,5210 | |
| | | | One sample Wilcoxon test (compared to 50) | | | CD: **p=0,0156**<br>VAD: **p=0,0156** | |
| Fig. S1B ♂ | Total distance (cm) over time (min) | CD: 9<br>VAD: 17 | 2-way RM ANOVA | Diet | F(1,23)=2,010 | p=0,1697 | |
| | | | | Time | F(1.597, 36.72)=17,70 | **p<0,0001** | |
| | | | | Interaction | F(9,207)=0,6565 | p=0,7475 | |
| Fig. S1B ♀ | Total distance (cm) over time (min) | CD: 7<br>VAD: 7 | 2-way RM ANOVA | Diet | F(1,12)=2,089 | p=0,1740 | |
| | | | | Time | F(3.242, 38.90)=22,51 | **p<0,0001** | |
| | | | | Interaction | F(9,108)=1,826 | p=0,0716 | |
| Fig. S1C ♂ | Total lever presses on RL (outcome devaluation) | CD: 7<br>VAD: 7 | 2-way ANOVA | Diet | F(1,24)=3,336 | p=0,0802 | |
| | | | | Devaluation | F(1,24)=112,5 | **p<0,0001** | |
| | | | | Interaction | F(1,24)=2,229 | p=0,1485 | |
| Fig. S1C ♀ | Total lever presses on RL (outcome devaluation) | CD: 7<br>VAD: 7 | 2-way ANOVA | Diet | F(1,24)=0,9369 | p=0,3427 | |
| | | | | Devaluation | F(1,24)=38,40 | **p<0,0001** | |
| | | | | Interaction | F(1,24)=0,9797 | p=0,3322 | |
| Fig. S1D ♂ | Total lever presses on UL (PRx2) | CD: 9<br>VAD: 15 | Normality test | Anderson-Darling | | **p=0,0033** | |
| | | | | D'Agostino-Pearson test | | **p=0,0021** | |
| | | | | Shapiro-Wilk | | **p=0,0011** | |
| | | | | Kolmogorov-Smirnov | | p>0,0001 | |
| | | | Mann-Whitney test | | | 0,0708 | |
| Fig. S1D ♀ | Total lever presses on UL (PRx2) | CD: 7<br>VAD: 7 | Normality test | Anderson-Darling | | p=0,2978 | |
| | | | | D'Agostino-Pearson test | | p=0,6314 | |
| | | | | Shapiro-Wilk | | p=0,6107 | |
| | | | | Kolmogorov-Smirnov | | p>0,0001 | |
| | | | Unpaired t-test | | | **p=0,0312** | |
| Fig. S1E ♂ | High reward lever preference | CD: 8<br>VAD: 9 | 2-way ANOVA | Diet | F(1,90)=15,74 | **p=0,0001** | |
| | | | | Delay | F(5,90)=101,7 | **p<0,0001** | |
| | | | | Interaction | F(5,90)=1,823 | p=0,1163 | |
| Fig. S1E ♀ | High reward lever preference | CD: 10<br>VAD: 10 | 2-way ANOVA | Diet | F(1,108)=1,437 | p=0,2332 | |
| | | | | Delay | F(5,108)=57,13) | **p<0,0001** | |
| | | | | Interaction | F(5,108)=0,7289 | p=0,6033 | |
| Fig. S1F ♂ | AUC of high reward lever preference | CD: 8<br>VAD: 9 | Normality test | Anderson-Darling | | p=0,3586 | |
| | | | | D'Agostino-Pearson test | | p=0,9788 | |
| | | | | Shapiro-Wilk | | p=0,4026 | |
| | | | | Kolmogorov-Smirnov | | p>0,0001 | |
| | | | Unpaired t-test | | | **p=0,0209** | |
| Fig. S1F ♀ | AUC of high reward lever preference | CD: 10<br>VAD: 10 | Normality test | Anderson-Darling | | p=0,1265 | |
| | | | | D'Agostino-Pearson test | | **p=0,0060** | |
| | | | | Shapiro-Wilk | | **p=0,0475** | |
| | | | | Kolmogorov-Smirnov | | p>0,0001 | |
| | | | Mann-Whitney test | | | p=0,5646 | |

| Figure panels | Outcome measure | n | Statistical analysis | | F-value | p-value | Multiple comparaisons |
|---|---|---|---|---|---|---|---|
| Fig. S2A ♂ | DS D1R relative expression (%) | CD: 23<br>VAD: 39 | Normality test | Anderson-Darling | | **p=0,0141** | |
| | | | | D'Agostino-Pearson test | | p=0,1781 | |
| | | | | Shapiro-Wilk | | **p=0,0305** | |
| | | | | Kolmogorov-Smirnov | | **p=0,0124** | |
| | | | Mann-Whitney test | | | p=0,2020 | |
| Fig. S2A ♀ | DS D1R relative expression (%) | CD: 20<br>VAD: 30 | Normality test | Anderson-Darling | | **p=0,0147** | |
| | | | | D'Agostino-Pearson test | | **p=0,0129** | |
| | | | | Shapiro-Wilk | | **p=0,0130** | |
| | | | | Kolmogorov-Smirnov | | **p=0,0085** | |
| | | | Mann-Whitney test | | | p=0,9884 | |
| Fig. S2B ♂ | DS D2R relative expression (%) | CD: 23<br>VAD: 39 | Normality test | Anderson-Darling | | p=0,0871 | |
| | | | | D'Agostino-Pearson test | | p=0,1507 | |
| | | | | Shapiro-Wilk | | **p=0,0261** | |
| | | | | Kolmogorov-Smirnov | | p>0,0001 | |
| | | | Mann-Whitney test | | | p=0,1673 | |
| Fig. S2B ♀ | DS D2R relative expression (%) | CD: 20<br>VAD: 30 | Normality test | Anderson-Darling | | **p=0,0179** | |
| | | | | D'Agostino-Pearson test | | **p=0,0005** | |
| | | | | Shapiro-Wilk | | **p=0,0061** | |
| | | | | Kolmogorov-Smirnov | | **p=0,0012** | |
| | | | Mann-Whitney test | | | p=0,9587 | |
| Fig. S2C ♂ | DS D3R relative expression (%) | CD: 23<br>VAD: 39 | Normality test | Anderson-Darling | | **p=0,0066** | |
| | | | | D'Agostino-Pearson test | | **p=0,0172** | |
| | | | | Shapiro-Wilk | | **p=0,0078** | |
| | | | | Kolmogorov-Smirnov | | **p=0,0521** | |
| | | | Mann-Whitney test | | | p=0,6322 | |
| Fig. S2C' ♀ | DS D3R relative expression (%) | CD: 20<br>VAD: 30 | Normality test | Anderson-Darling | | **p=0,0144** | |
| | | | | D'Agostino-Pearson test | | **p=0,0020** | |
| | | | | Shapiro-Wilk | | **p=0,0044** | |
| | | | | Kolmogorov-Smirnov | | **p=0,0355** | |
| | | | Mann-Whitney test | | | p=0,0601 | |
| Fig. S2D ♂ | VTA TH relative expression (%) | CD: 14<br>VAD: 27 | Normality test | Anderson-Darling | | p=0,5828 | |
| | | | | D'Agostino-Pearson test | | p=0,4411 | |
| | | | | Shapiro-Wilk | | p=0,3344 | |
| | | | | Kolmogorov-Smirnov | | p>0,0001 | |
| | | | Unpaired t-test | | | p=0,2163 | |
| Fig. S2D ♀ | VTA TH relative expression (%) | CD: 15<br>VAD: 22 | Normality test | Anderson-Darling | | p=0,8395 | |
| | | | | D'Agostino-Pearson test | | p=0,6558 | |
| | | | | Shapiro-Wilk | | p=0,6686 | |
| | | | | Kolmogorov-Smirnov | | p>0,0001 | |
| | | | Unpaired t-test | | | p=0,5332 | |
| | | | | Anderson-Darling | | p=0,4742 | |

| | | | | | | |
|---|---|---|---|---|---|---|
| Fig. S2E ♂ | VTA DDC relative expression (%) | CD: 14<br>VAD: 27 | Normality test | D'Agostino-Pearson test | | p=0,9448 |
| | | | | Shapiro-Wilk | | p=0,6444 |
| | | | | Kolmogorov-Smirnov | | p>0,0001 |
| | | | Unpaired t-test | | | p=0,2495 |
| Fig. S2E ♀ | VTA DDC relative expression (%) | CD: 15<br>VAD: 22 | Normality test | Anderson-Darling | | p=0,8921 |
| | | | | D'Agostino-Pearson test | | p=0,7811 |
| | | | | Shapiro-Wilk | | p=0,9326 |
| | | | | Kolmogorov-Smirnov | | p>0,0001 |
| | | | Unpaired t-test | | | p=0,2719 |
| Fig. S2F ♂ | VTA MAOA relative expression (%) | CD: 14<br>VAD: 27 | Normality test | Anderson-Darling | | p=0,0509 |
| | | | | D'Agostino-Pearson test | | p=0,1691 |
| | | | | Shapiro-Wilk | | p=0,1031 |
| | | | | Kolmogorov-Smirnov | | **p=0,0321** |
| | | | Mann-Whitney test | | | p=0,8599 |
| Fig. S2F ♀ | VTA MAOA relative expression (%) | CD: 15<br>VAD: 22 | Normality test | Anderson-Darling | | p=0,1282 |
| | | | | D'Agostino-Pearson test | | p=0,2733 |
| | | | | Shapiro-Wilk | | p=0,2493 |
| | | | | Kolmogorov-Smirnov | | p=0,0984 |
| | | | Unpaired t-test | | | p=0,1011 |
| Fig. S2G ♂ | VTA VMAT relative expression (%) | CD: 14<br>VAD: 27 | Normality test | Anderson-Darling | | p=0,5202 |
| | | | | D'Agostino-Pearson test | | p=0,7887 |
| | | | | Shapiro-Wilk | | p=0,8295 |
| | | | | Kolmogorov-Smirnov | | p>0,0001 |
| | | | Unpaired t-test | | | p=0,1206 |
| Fig. S2G ♀ | VTA VMAT relative expression (%) | CD: 15<br>VAD: 22 | Normality test | Anderson-Darling | | **p=0,0494** |
| | | | | D'Agostino-Pearson test | | **p=0,0029** |
| | | | | Shapiro-Wilk | | **p=0,0136** |
| | | | | Kolmogorov-Smirnov | | p>0,0001 |
| | | | Mann-Whitney test | | | p=0,6211 |
| Fig. S2H | VTA RARβ mRNA expression / VTA DAT mRNA expression correlation | CD: 9<br>VAD: 14 | Pearson correlation | | | $R^2$=0.2552<br>**p=0.0139** |
| Fig. S2J | Number of TH+ neurons | CD: 9<br>VAD: 4-6 | 2-way ANOVA | Diet | F(1,24)=0,02118 | p=0,8855 |
| | | | | Structure | F(1,24)=3,588 | p=0,0703 |
| | | | | Interaction | F(1,24)=0,4743 | p=0,4976 |
| Fig. S2L | TH immunostaining intensity | CD: 7<br>VAD: 6 | 2-way ANOVA | Diet | F(1,55)=1,194 | p=0,2792 |
| | | | | Structure | F(4,55)=1,768 | p=0,1485 |
| | | | | Interaction | F(4,55)=0,5885 | p=0,6723 |

| Figure panels | Outcome measure | n | Statistical analysis | | F-value | p-value | Multiple comparaisons |
|---|---|---|---|---|---|---|---|
| Fig. S3B | Lick z-score in FR1/FR5/FR10 | CD: 5<br>VAD: 7 | 2-way ANOVA | Diet | F(1,28)=0,07797 | p=0,7821 | |
| | | | | Ratio | F(2,28)=3,369 | **p=0,0488** | |
| | | | | Interaction | F(2,28)=0,7197 | p=0,7821 | |
| Fig. S3D | UL press in FR1/FR5/FR10 | CD: 5<br>VAD: 7 | 2-way ANOVA | Diet | F(1,29)=11,09 | **p=0,0024** | |
| | | | | Ratio | F(2,29)=0,4890 | p=0,6182 | |
| | | | | Interaction | F(2,29)=0,07274 | p=0,9300 | |
| Fig. 3E | Number of licks (Pavlovian) | CD: 10<br>VAD: 11 | Normality test | Anderson-Darling | | p=0,1597 | |
| | | | | D'Agostino-Pearson test | | **p=0,0215** | |
| | | | | Shapiro-Wilk | | p=0,1142 | |
| | | | | Kolmogorov-Smirnov | | p>0,1000 | |
| | | | Mann-Whitney test | | | p=0,4078 | |
| Fig. 3E' | Number of reward (Pavlovian) | CD: 10<br>VAD: 11 | Normality test | Anderson-Darling | | **p=0,0133** | |
| | | | | D'Agostino-Pearson test | | **p=0,0072** | |
| | | | | Shapiro-Wilk | | **p=0,0141** | |
| | | | | Kolmogorov-Smirnov | | **p=0,0036** | |
| | | | Mann-Whitney test | | | p=0,6970 | |
| Fig. S3F | Number of licks (FR1, FR5 and FR10) | CD: 5<br>VAD: 7 | 2-way ANOVA | Diet | F(1,30)=1,039 | p=0,3162 | |
| | | | | Ratio | F(2,30)=1,653 | p=0,2085 | |
| | | | | Interaction | F(2,30)=0,08997 | p=0,9142 | |
| Fig. S3F' | Number of reward (FR1, FR5 and FR10) | CD: 5<br>VAD: 7 | 2-way ANOVA | Diet | F(1,30)=1,464 | p=0,2357 | |
| | | | | Ratio | F(2,30)=3,066 | p=0,0614 | |
| | | | | Interaction | F(2,30)=0,1978 | p=0,8216 | |
| Fig. S3F'' | Total lever presses on reinforced lever (FR1, FR5 and FR10) | CD: 5<br>VAD: 7 | 2-way ANOVA | Diet | F(1,30)=0,2487 | p=0,6216 | |
| | | | | Ratio | F(2,30)=21,20 | **p<0,0001** | |
| | | | | Interaction | F(2,30)=0,1715 | p=0,8432 | |
| Fig. S3F''' | Total lever presses on non-reinforced lever (FR1, FR5 and FR10) | CD: 5<br>VAD: 7 | 2-way ANOVA | Diet | F(1,30)=0,4577 | p=0,5039 | |
| | | | | Ratio | F(2,30)=3,438 | **p=0,0452** | |
| | | | | Interaction | F(2,30)=2,631 | p=0,0886 | |

| Figure panels | Outcome measure | n | Statistical analysis | | F-value | p-value | Multiple comparaisons |
|---|---|---|---|---|---|---|---|
| Fig. S4A | Total lever presses on RL in CD mice saline vs CNO | 9 | Normality test | Anderson-Darling | | p=0,189 | |
| | | | | D'Agostino-Pearson test | | p=0,0858 | |
| | | | | Shapiro-Wilk | | p=0,2262 | |
| | | | | Kolmogorov-Smirnov | | p>0,1000 | |
| | | | Unpaired t-test | | | **p=0,0171** | |
| Fig. S4A | Total lever presses on RL in VAD mice saline vs CNO | 10 | Normality test | Anderson-Darling | | p=0,6631 | |
| | | | | D'Agostino-Pearson test | | p=0,814 | |
| | | | | Shapiro-Wilk | | p=0,8009 | |
| | | | | Kolmogorov-Smirnov | | p>0,1000 | |
| | | | Unpaired t-test | | | p=0,2136 | |
| Fig. 4E | Total lever presses on RL (concurrent task) under CNO | mcherry CD : 8<br>mcherry VAD: 8<br>Gi CD: 8<br>Gi VAD: 8 | 2-way ANOVA | Diet | F(1,28)=0,3699 | p=0,5340 | |
| | | | | Virus | F(1,28)=3,425 | p=0,0748 | |
| | | | | Interaction | F(1,28)=0,04034 | p=0,8123 | |
| Fig. 4F | Chow consumption (concurrent task) under CNO | mcherry CD : 8<br>mcherry VAD: 8<br>Gi CD: 8<br>Gi VAD: 8 | 2-way ANOVA | Diet | F(1,28)=0,03705 | p=0,8488 | |
| | | | | Virus | F(1,28)=11 | **p=0,0025** | |
| | | | | Interaction | F(1,28)=0,2943 | p=0,5918 | |
